# Combinatorial cooperativity can facilitate epithelial-mesenchymal transition in a miR200-Zeb feedback network

**DOI:** 10.1101/2023.10.06.561220

**Authors:** Mubasher Rashid, Brasanna M Devi, Malay Banerjee

## Abstract

Carcinoma cells often utilize epithelial-mesenchymal transition (EMT) programs for cancer progression and metastasis. Numerous studies point to the SNAIL-induced miR200/Zeb feedback circuit as crucial in regulating EMT by shifting cancer cells to at least three (epithelial (E), hybrid (h E/M), mesenchymal (M)) phenotypic states arrayed along the epithelial-mesenchymal phenotypic spectrum. However, a coherent molecular-level understanding of how such a tiny circuit controls carcinoma cell entrance into and residence in various states is lacking. Here, we use molecular binding data and mathematical modeling to report that miR200/Zeb circuit can essentially utilize combinatorial cooperativity to control E-M phenotypic plasticity. We identify minimal combinatorial cooperativities that give rise to E, hybrid-E/M, and M phenotypes. We show that disrupting a specific number of miR200 binding sites on Zeb as well as Zeb binding sites on miR200 can have phenotypic consequences – the circuit can dynamically switch between two (E, M) and three (E, hybrid-E/M, M) phenotypes. Further, we report that in both SNAIL-induced and SNAIL knock-out miR200/Zeb circuits, cooperative transcriptional feedback on Zeb as well as Zeb translational inhibition due to miR200 are essential for the occurrence of intermediate hybrid-E/M phenotype. Finally, we demonstrate that SNAIL can be dispensable for EMT, and in the absence of SNAIL, the transcriptional feedback can control cell state transition from E to hybrid-E/M, to M state. Our results thus highlight molecular-level regulation of EMT in miR200/Zeb circuit and we expect these findings to be crucial to future efforts aiming to prevent EMT-facilitated dissemination of carcinomas.

## Introduction

Despite significant advances in diagnosing and treating cancer, metastasis persists as a barrier to successful therapy and is the main cause of cancer-related death^1,2^. The early steps of this highly complex process rely on non-motile epithelial tumor cells acquiring characteristics of mesenchymal cells^3^, also called as epithelial-to-mesenchymal transition, which are more migratory^4,5^. The migrating cancer cells then undergo a reverse mesenchymal-to-epithelial transition (MET) to seed metastatic tumors^3^. Accumulating evidence has demonstrated that induction of EMT facilitates carcinoma cell dissemination^6,7^, entrance into stem-cell-like states^8,9^, and resistance to cell death induced through various therapeutic treatments^10–13^. By enabling carcinoma cell phenotypic plasticity^14,15^, an EMT program can shift cancer cells to various phenotypic states arrayed along the epithelial (E) to mesenchymal (M) spectrum^16–18^, leading to phenotypic diversification within individual tumours, also referred to as non-genetic intra-tumor heterogeneity^19^. Recent reports have also emphasized that tumor progression and metastasis requires EMT to be transient and reversible, giving rise to continuum of intermediate states. Carcinoma cells residing in intermediate phenotype, also known as hybrid epithelial-mesenchymal (h-E/M) state, are associated with malignant characteristics, such as invasiveness, tumor-initiating ability, therapy resistance, and metastatic dissemination^17,20–22^. However, understanding the molecular controls that enable carcinoma cells to enter and dwell in one or another phenotypic state along the E–M spectrum is still a major challenge^23^. Identifying these controls can be one of the first steps to design effective therapeutics to arrest phenotypic plasticity of carcinoma cells and prevent metastasis.

With the advent of high-throughput molecular profiling techniques, network-based models and approaches have proved extremely useful to understand cell phenotypic transition across contexts^24,25^. In the case of EMT, particularly, computational models of transcriptional networks (including transcription factors, micro-RNAs, mRNAs, etc.) underlying EMT have associated differential expression patterns of specific molecules to h-E/M phenotype^26–29^. Some more recent studies have associated network characteristics such as network-frustration^30^, besides other topological features of large and complex EMT networks to the emergence of h-E/M phenotype ^31^, as well as proposed inhibitors of E-M plasticity^32^. Nevertheless, these EMT transcriptional networks studied for carcinoma cell-fate transition and plasticity are large and complex^31^. However, interestingly, a common building block of these networks is miR200/Zeb network motif; in which miR200 microRNA - an epithelial marker, and Zeb mRNA – a mesenchymal marker, mutually inhibit each other to form a microRNA-mRNA toggle switch^27,33–36^ **(Fig. 1)**. This circuit integrated with other EMT molecular networks is proposed to be a “*motor of cellular plasticity*”^37^, besides regulating stemness during the generation of cancer stem cells^38^, tumor differentiation and invasion^39^, and controlling key cell-cell communication pathway involving Notch/Jagged1^40–42^. Due to such a diverse functionality, this little circuit, as part of relatively bigger networks, has been widely studied for EMT as well, in different carcinomas^21,28,43^. A natural question that we wanted to ask is: whether miR200/Zeb, as an independent network, can recapitulate the three phenotypes – E, M and h-E/M, observed in relatively larger networks of which this network is a part? Answering this question may reveal additional independent, and perhaps surprising, roles of this circuit, in E-M plasticity and can narrow down efforts focussing on large networks to prevent EMT and metastasis.

A glimpse at the network-topology of miR200/Zeb circuit shows that it is not a complex circuit, since it just has two molecules with few interactions in between. However, a closer look at the molecular interaction details reveals inherent complexity in terms of the number as well as different type of binding sites on each molecule. The miR200 harbours three sites to bind Zeb and two sites to bind SNAIL, while Zeb harbours six/three sites to bind miR200a/miR200b,c, and two sites to bind SNAIL, besides two more sites to bind its own transcription factor, ZEB. Further details are in the scheme below. We refer to this type of mechanism, where several molecules, each with positive cooperativity, combinatorially regulate a particular molecule, as *combinatorially cooperative (CC)*. A question that remains unanswered until now is, can a particular choice of CC decide the phenotypic landscape outcome of miR200/Zeb circuit, so that a link can be established between binding cooperativity and phenotypic heterogeneity. Addressing this question can provide molecular level explanation to phenotypic plasticity enabled by miR200/Zeb circuit.

Using maximal CC (i.e., all binding sites on every molecule are occupied), *Lu et al.,*^44^ demonstrated SNAIL driven miR200a/Zeb circuit can act as a ternary switch, giving rise to h-E/M phenotype, besides E and M phenotypes. We then ask, whether tristability can be achieved in miR200a/Zeb circuit in which miR200b,c can bind to just three sites on Zeb, or in a generic case, when only a few binding sites on each molecule are occupied. And, if so, what could be the minimum such number for each molecule?

To answer this question, we analysed the steady state dynamics of SNAIL driven miR-200a/ZEB and miR200a/Zeb circuits and mapped the stable steady states to specific phenotypes based on the known E and M markers. We report that maximal CC is not required, in either circuits, to generate three phenotypes, and identified least number of binding sites that must be occupied to observe tristability. We also obtained necessary conditions to observe three phenotypes: (1) Zeb mRNA self-activation must have a cooperativity of order at least two. (2) miR200 to Zeb mRNA binding must have a cooperativity of order at least three. Further, we refute the assertion by Lu et al.,^44^ and demonstrate that simultaneous active degradation and translation inhibition of Zeb by miR200 are not required to attain h-E/M phenotype and that translation inhibition alone is sufficient. Further, we show that even SNAIL induction on miR200/Zeb circuits may not be required for EMT and that in the absence of SNAIL, the strength of self-feedback loop on Zeb can determine the phenotypic transition – low to high levels drive cells out from E to h-E/M to M states. This result is partially in line with a recent *in vivo* study on mouse model of pancreatic ductal adenocarcinomas showing that SNAIL deletion does not influence the metastatic phenotype and is thus not rate limiting for EMT and metastasis^10^. Our results thus unreveal the previously unknown role of combinatorial cooperative binding in EMT and uphold that by manipulating CC, miR200/Zeb axis can independently act as a motor of cellular heterogenity. These results corraborate with the findings of Title et al.,^39^ which report, in two epithelial cancer models, that mutating the miR200 binding sites on Zeb are sufficient to induce EMT. Our findings thus suggest that future clinical efforts to arrest phenotypic plasticity enabled by EMT should focus on molecular interactions between miR200 and Zeb rather than targeting complex EMT networks.

## Results

### Schematic of miR200/Zeb circuit

Before presenting our results, we will discuss the details of interactions between the molecules in SNAIL induced miR200/Zeb circuit represented by the schematic in **Fig. 1**. At the transcription level, miR200 harbours two and three sites to bind SNAIL and Zeb-TF (here onwards referred as ZEB), respectively^45,46^. Also, Zeb harbours four sites, two each to bind SNAIL and ZEB, and activate its expression^4046^. On the other, at the translation level, Zeb harbours six/three sites to bind miR200a/miR200b,c molecules and inhibit or degrade Zeb^39^. Moreover, unlike classical TF-TF toggle switch in which molecules interact at transcription level, microRNA-mRNA circuit can additionally involve three types of interactions at the translation level. Firstly, miR200 can directly bind to Zeb and degrade it, referred to as active degradation of Zeb. Secondly, miR200 can directly bind to Zeb and prevent it from translating into a ZEB protein, referred to as translation inhibition of Zeb. Finally, during the process of miR200-Zeb complex formation, miR200 itself poses a chance of being degraded, which is referred to as active degradation of miR200^47^. As shown in the scheme, both miR200 and Zeb-mRNA molecules are combinatorially regulated by at least two other molecules with cooperativity greater or equal to two. This mechanism of regulation of a molecule by multiple molecules with higher order cooperativity is termed as combinatorial cooperativity.

**Fig. 1.**
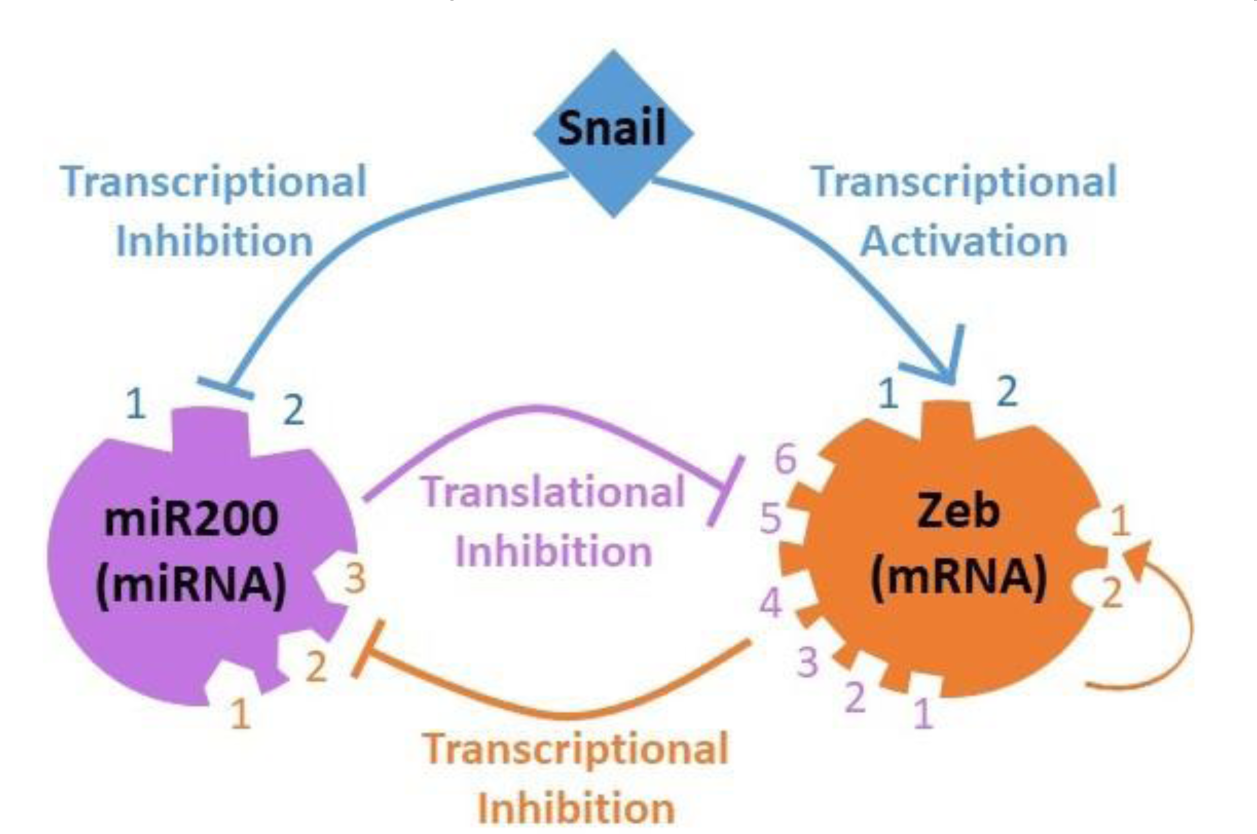
Scheme of molecular interactions in miR200/Zeb network. Both miR200 (purple) and Zeb (orange) inhibit expression of each other. The miR200 collectively represents both miR200a and miR200b,c family members. SNAIL (blue) acts on the top of miR200/Zeb circuit by repression miR200 and activating Zeb. The self-loop on Zeb shows that Zeb can activate its own expression. Numbers represent binding-site count on each molecule. Curves with arrowheads denote activation and those with bars denote inhibition or repression.

### Mathematical modeling Framework

In this section, we will describe the framework to model translation inhibition and active degradation as well as transcription activation/inhibition. As discussed in the scheme, miR200 can form a complex with Zeb that results in silencing of Zeb through translation inhibition (purple edge in **Fig. 1**) and active degradation. In this process, miR200 also poses a chance to be actively degraded. To explicitly include the role of each miR200-Zeb complex in active degradation and translation inhibition, we use the following modeling framework adopted from Lu et al.,^43^.

### Modeling translation inhibition and active degradation

An mRNA having *n* binding sites for miRNA (notation for micro-RNA), can exist in *n* + 1 possible states – one native state (with no miRNA bound) and *i* = 1, 2, . . ., *n* miRNA bound states. We make an assumption here that binding/unbinding rates of miRNA and mRNA are fast, compared to their production and degradation rates and binding is independent for different binding sites. Denoting the concentrations of miRNA and mRNA by [*X*] and [*Y*], resp., it follows that, at equilibrium, the concentration of mRNA with *i* miRNA bound (denoted by [Y]^*i*^) obeys,

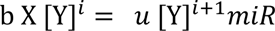

So that,

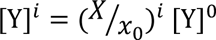

Here, b and u represent binding and unbinding rates, resp., [Y]^0^ denotes mRNA with no miRNA bound to it, and 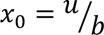 is the unbinding to binding ratio.

The total concentration of mRNA (*Y*) then satisfies,

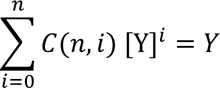

Where, 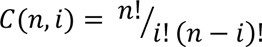 denotes the configuration when *i* out of *n* miRNA’s are bound to mRNA. So that,

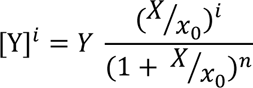

Thus, the net active degradations of *X*, *Y*, and net translation inhibition of *Y* are given by equations *M*_1_, *M*_2_, and *M*_3_, respectively.

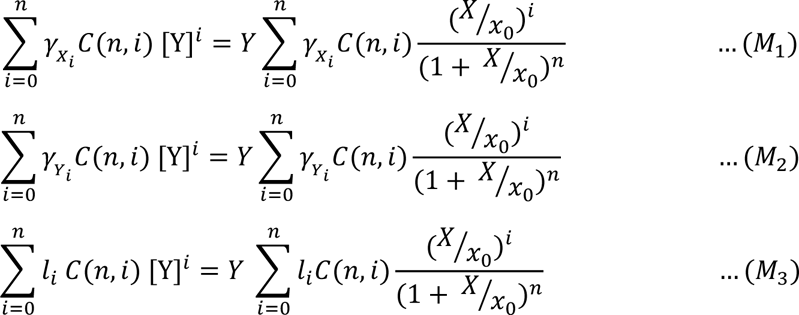

Where, *γ_X_i__* and *γ_Y_i__* denote active degradation rates of *X* and *Y*, resp., and *l*_*i*_ denotes translation inhibition rate of *Y*, when *i* out of *n* sites are occupied.

### Modeling transcription activation/inhibition

Transcription interactions (blue and yellow edges in **Fig. 1**) are commonly modeled using Hill functions. We however use a shifted Hill function defined in *M*_4_to take into consideration the fold change (*λ*) caused in the basal expression of molecule *A* (say) due to transcription activation/inhibition by molecule *B* (say).

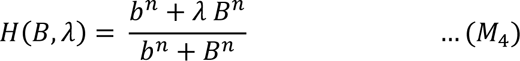

So that,

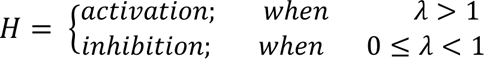

Here, *n* is the Hill coefficient and denotes binding cooperativity.

### Model of SNAIL induced miR200/Zeb network

By adopting the modeling framework discussed in equations *M*_1_ − *M*_4_, the system of coupled non-linear ordinary differential equations in (1) describe the dynamics of the network in **Fig. 1**.

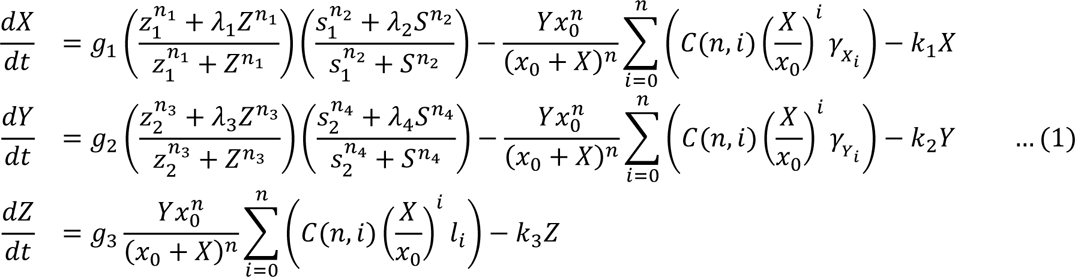

Where, *g*_*i*_, *i* = 1, 2, 3, resp., represents synthesis rates of miR200, Zeb, and ZEB. *z*_*i*_, *i* = 1, 2, resp., represent ZEB binding thresholds to miR200 and Zeb. *λ*_*i*_, *i* = 1, 2, resp., represent SNAIL binding threshold to miR200 and Zeb. *λ*_*i*_, *i* = 1, 2, 3, 4, resp., represent fold change in transcription rate due to ZEB binding to miR200, SNAIL binding to miR200, ZEB binding to Zeb, and SNAIL binding to Zeb. *k*_*i*_, *i* = 1,2,3, resp, represent innate degradation rates of miR200, Zeb, and ZEB. *x*_0_ is the ratio of unbinding to binding rates of miR200 to Zeb. *n*_*i*_, *i* = 1, 2, 3, 4, resp., are the Hill coefficient bindings (or transcriptional binding cooperativities) of ZEB to miR200, SNAIL to miR200, ZEB to Zeb, and SNAIL to Zeb. Note that *n* represents translational cooperativity of miR200 binding to Zeb, so that that *n* = 6 mimics miR200a/Zeb circuit and *n* = 3 mimics miR200b,c/Zeb circuit. *S* denotes SNAIL concentration. The parameters for model (1) are adopted from Lu et al^44^ who have curated them after taking into account all possible experimental data. Interestingly, most of these parameters work for models (2), (3), and (4) as well, with slight changes in a few. Parameter values are reported in **Table S1, S2** in *Supplementary material*.

In what follows, is the investigation of EMT and tristability (particularly focusing on h-E/M phenotype), by considering four versions of model (1) under different CC conditions: (i) In model (1), we consider both translation inhibition (TI) and active degradation (AD). (ii) In model (2), we investigate whether active degradations of miR200 and Zeb are really required to attain tristability; we thus drop active degradations in this version and study the circuits with only translation inhibition. (iii) In model (3), we investigate whether EMT can happen without the induction of SNAIL and we thus consider a circuit with both TI and AD, but without SNAIL. (iv) In model (4), we investigate whether miR200/Zeb alone can exhibit tristability so that we consider a circuit without SNAIL and AD’s of miR200b,c and Zeb. In each model, except SNAIL knock-out ones, we also report minimum values of CC, i.e., lower bound for 5-tuple, (*n*; *n*_1_, *n*_2_, *n*_3_, *n*_4_), that can support E, M and h-E/M phenotypes. In SNAIL knock-in models, however, we report lower bound for 3-tuple, (*n*; *n*_1_, *n*_3_), that can support E, M and h-E/M phenotypes.

### Tristability in miR200a/Zeb and miR200b,c/Zeb circuits with both active degradation and translation inhibition

Recent theoretical and experimental studies reported tri-stability (E, M and h-E/M phenotypes) in SNAIL driven miR200a/Zeb circuit in which miR200a binds to six sites on Zeb, Zeb binds to three sites on miR200a, SNAIL binds to two sites each on miR200 and Zeb, and ZEB binds to two sites on Zeb Zeb^44,48^. However as reported in Burk et al.,^45^ miR200b,c family of microRNA’s bind only up to three conserved binding sites on Zeb. We thus ask whether three miR200b,c binding sites on Zeb can have phenotypic consequences in EMT. And if so, what is the functional role of six miR200a binding sites on Zeb compared to just three miR200b,c binding sites.

To test this hypothesis, we first recapitulated the three phenotypes (E, M and h-E/M states) observed in Lu et al.,^44^ by mimicking their experiment with (*n* = 6; *n*_1_ = 3, *n*_2_ = 2, *n*_3_ = 2, *n*_4_ = 2) in model (1). As shown in phase plane in **Fig. 2A**, the three stable states given by (high miR200, low Zeb), (partial miR200, partial Zeb), and (low miR200, high Zeb) exist concurrently. Denoting the low, intermediate, and high levels of a molecule by 0, ½, and 1, resp., the three states can then be represented as (1, 0), (½, ½), and (0, 1). Since miR200 is a known epithelial marker and Zeb a mesenchymal marker, we designate the three stable states as E, h-E/M, and M phenotypes. Next, we mimicked the miR200b,c/Zeb circuit by changing (*n* = 3) in model (1) while keeping all other parameters in Table S1, S2, unchanged. As shown in the nullcline plot in **Fig. 2B** shows, we were again able to achieve the three phenotypes thereby supporting our hypothesis that the three miR200b,c sites on Zeb are sufficient to achieve tri-stability.

**Fig. 2.**
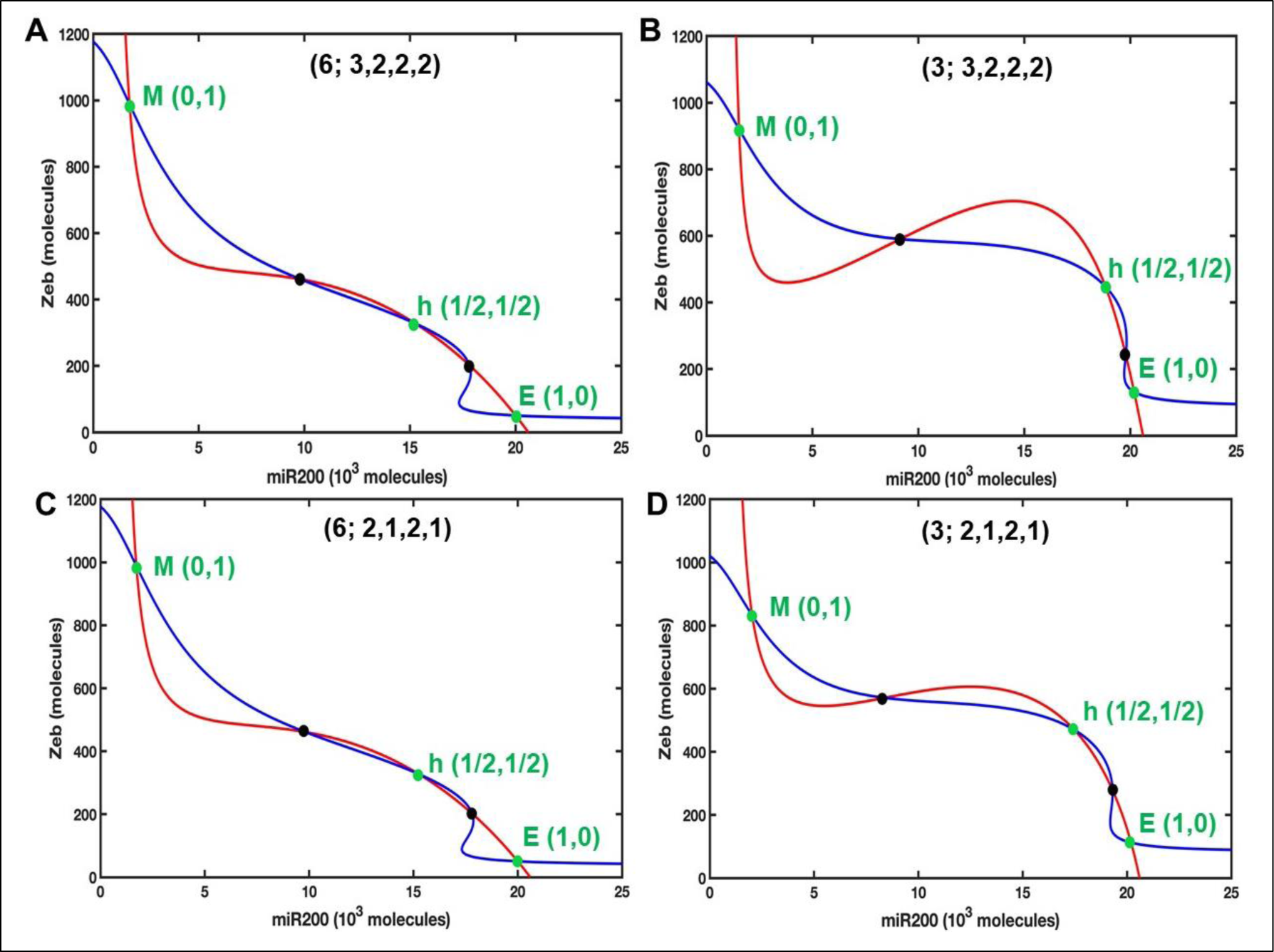
Tri-stability in miR200/Zeb circuit. Nullcline plots showing existence of three stable states corresponding E, h-E/M and M phenotypes in miR200/Zeb circuits. The five intersections of the curves correspond to five steady states. Green dots represent stable steady states and black dots represent unstable steady states. Tri-stability in miR200a/Zeb (A) and miR200b,c/Zeb (B) circuits with maximum CC. Tristability in the minimal CC counterparts is shown in (C) and (D), resp. Red/blue nullcline is obtained by substituting third equation in first and second equations and then setting first/second equation in model (1) to zero. The miR200 is E marker so that low/high levels of miR200 correspond to E/M state. Similarly, Zeb is M marker so that low/high levels of Zeb correspond to M/E states. Intermediated levels of these molecules correspond to h-E/M states.

To identify the minimal CC thresholds that are sufficient to exhibit the three phenotypes observed in relatively highly non-linear circuits, we tried different combinations of (*n*_1_, *n*_2_, *n*_3_, *n*_4_). Interestingly, we found that in both miR200a/Zeb (*mimicked by n* = 6) (**Fig. 2C**) and miR200b,c/Zeb (*mimicked by n* = 3) (**Fig. 2C**) circuits, the minimal CC that preserves the three phenotypes is (*n*_1_ = 2, *n*_2_ = 1, *n*_3_ = 2, *n*_4_ = 1). Since *n*_1_ and *n*_3_ mimic the binding cooperativity of ZEB for miR200 and Zeb itself (self-loop), resp., we conclude that cooperative self-activation of Zeb as well as cooperative inhibition of miR200 by Zeb is essential to achieve tri-stability in either circuits.

### EMT in miR200a/Zeb and miR200b,c/Zeb circuits under different CC conditions

Experimental studies have reported SNAIL as one of the drivers of cell state transition during EMT in different types of carcinomas. The SNAIL induced EMT has been observed in miR200a/Zeb circuit where induction of SNAIL causes E cells to switch to h-E/M state and further to M state^44^. Whether miR200/Zeb circuits can also exhibits EMT upon SNAIL induction and more importantly, whether cooperativity has any role in the stability of phenotypes is unknown. To address this question, we performed a comparative analysis of SNAIL induced EMT in miR200a/Zeb and miR200b,c/Zeb circuits using different choices of CC. Bifurcation diagrams illustrating the coexistence of three states and the transition in between are presented in **Fig. 3**. In miR200a/Zeb circuit **E** phenotype switches to **h-E/M** phenotype that further switches to **M** phenotype for increased SNAIL levels (**Fig. 3A**). A similar dynamics is observed in miR200b,c/Zeb circuit, though the phenotypic transition happens for relatively low levels of SNAIL (**Fig. 3B**). In both circuits, it is evident that while **E** to **M** transition occurs via an **h-E/M state**, **M** to **E** transition happens directly without any intermediate state. These data corroborate with the experimental findings reporting the non-transitioning to hybrid phenotype during MET^49,50^.

**Fig. 3.**
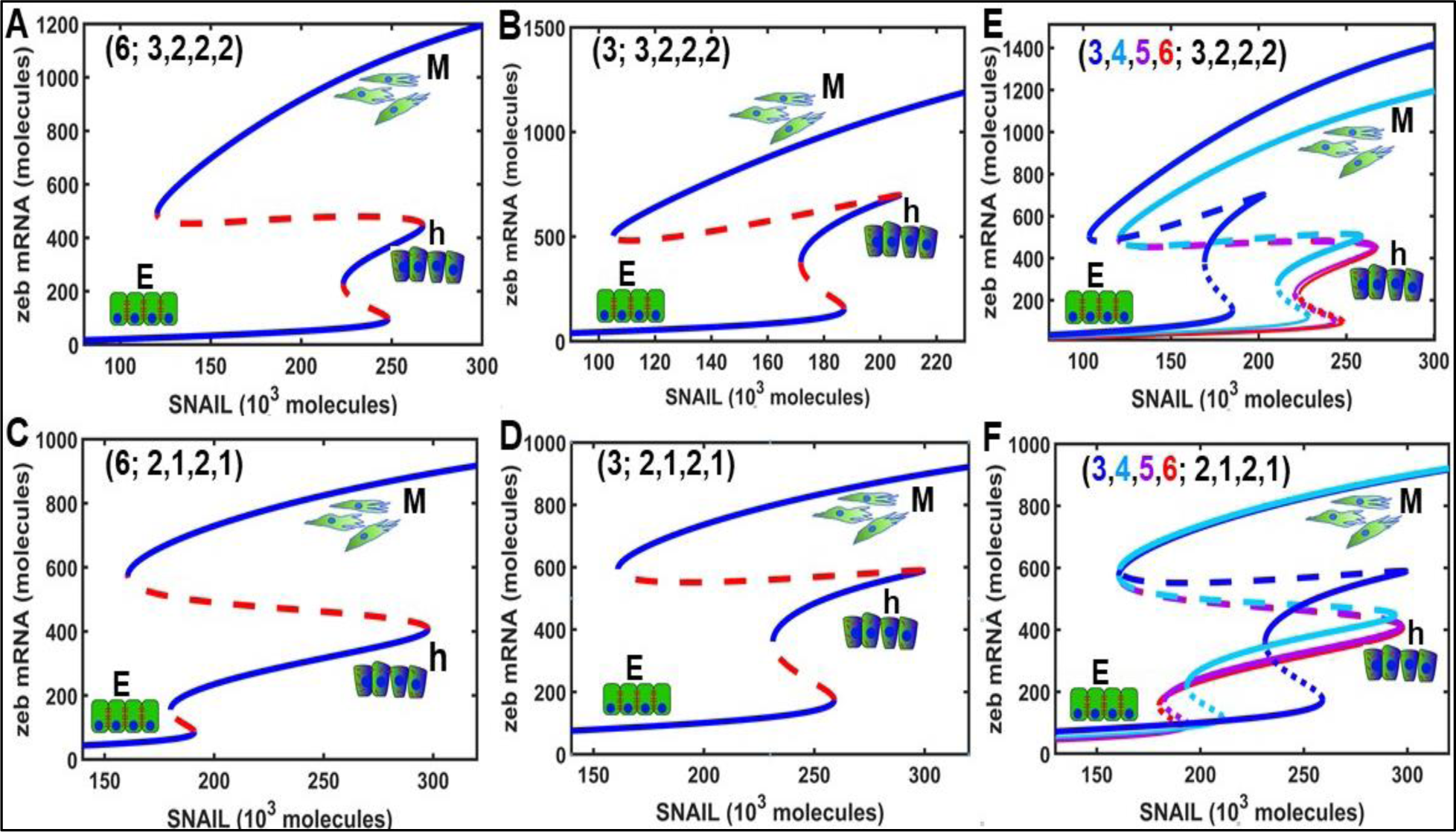
Bifurcation diagrams showing EMT under different cooperativity conditions. Low Zeb mRNA and SNAIL levels correspond to E phenotype, intermediate levels represent h-E/M phenotype and high levels represent M phenotype. First vertical panel represents miR200a/Zeb circuit with higher (A) and lower (C) order cooperativity. Second vertical panel represents miR200b,c/Zeb circuit with higher (B) and lower (D) order cooperativity. In third vertical panel, as n varies from three to six, the circuit gradually shifts from highly cooperative miR200b,c/Zeb circuit (E, blue curve) to highly cooperative miR200a/Zeb circuit (E, red curve) and from the least cooperative miR200b,c/Zeb circuit (F, blue curve) to the least cooperative miR200a/Zeb circuit (F, red curve). With low SNAIL levels, Zeb levels remain low which correspond to E phenotype. With increased SNAIL levels, EMT proceeds via h-E/M phenotype. However, a direct transition from M to E is observed with decreased SNAIL levels. Solid and dotted curves represent stable and unstable steady states, respectively.

We further asked whether the least cooperative (weakly non-linear) versions of either circuits can reproduce tri-stability as well as explain transition between the three phenotypes. We tried different sets of CC and like earlier we found that (2, 1, 2, 1) combination in either circuits can produce similar dynamics as their highly cooperative (highly non-linear) “wild type” counterparts. We, however, found that in the least cooperative miR200a/Zeb circuit, though the three phenotypes coexist for relatively narrow range of SNAIL (**Fig. 3C**) compared to its highly cooperative counterpart (**Fig. 3A**), the **h-E/M phenotype** is more stabilized in the former circuit. In miR200b,c/Zeb circuit, however, both the highly cooperative (**Fig. 3B**) as well as the least cooperative circuits (**Fig. 3D**) show similar dynamics. A cross comparison of the four circuits shows that the best dynamics – in terms of the stability of **h-E/M phenotype** and the range of SNAIL levels for which the three states coexist, is observed in a least cooperative miR200b,c/Zeb circuit. We thus conclude that “matched or optimal” cooperativity at the transcription and translation levels, that’s achieved in least cooperative miR200b,c/Zeb circuit, might be one of the deciding factors in the emergence of **h-E/M phenotype** and therefore in phenotypic heterogeneity during EMT and metastasis.

Next, we varied posttranslational cooperativity of miR200 binding to Zeb between three to six in both the circuits and analysed EMT. We found that in highly cooperative circuit, as we move from third to sixth degree binding cooperativity, the tri-stable region stretches gradually (**Fig. 3E**). In contrast, an opposite behaviour is observed in least cooperative circuit where reduction in cooperativity at posttranslational level stretches the tri-stable region as well as increases the stability of the h-E/M phenotype (**Fig. 3F**). This analyses further confirms the role of matching cooperativity at transcription and posttranslational levels in the stability of intermediate h-E/M phenotype.

### Active degradation of Zeb due to miR200 may not be required for tristability

As discuss earlier, micro-RNA’s silence mRNA’s through active degradation and/or translation inhibition, with later more prominent than former. These silencing mechanisms can differentially affect the steady state dynamics of the micro-RNA involved circuits. We asked whether simultaneous activation of these mechanisms is required for tristability or each can independently work to achieve tristability. To investigate this, we decoupled the two mechanisms modelled in (1) by dropping (unaccounted for) active degradation and obtained model (2) with only translation inhibition. The translation inhibition of Zeb due to miR200 by binding to its six sites is modelled using summation in the third equation in (2). So that, n=6 refers to miR200a/Zeb circuit, while n=3 refers to miR200b,c/Zeb circuit.

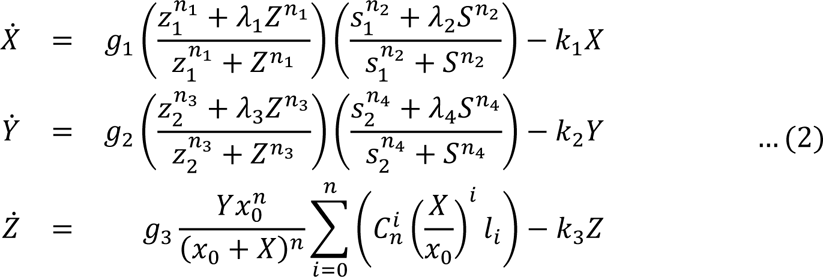

Lu et al.,^44^ suggested that absence of miR200 assisted active degradation in a highly cooperative miR200a/Zeb circuit may result into bistable dynamics. We, however, refute their assertion by showing that active degradation is not required to attain three stable states, i.e., tristability. As shown in the nullcline plot the three stable states exist in highly cooperative miR200a/Zeb circuit (**Fig. S1A**) as well as in highly cooperative miR200b,c/Zeb circuit (**Fig. S1B**). We further report that the three stable states are possible in least cooperative versions of miR200a/Zeb (**Fig. S1C**) and miR200b,c/Zeb (**Fig. S1D**) circuits as well, though the states are only best characterized in their highly cooperative counterparts.

To investigate the stability of the states and the transition in between, we found that with the continuous induction of SNAIL on highly cooperative miR200a/Zeb circuit, **E** to **M** transition happens through **h-E/M state** while the reverse (M to E) happens directly (**Fig. 4A**). The three state (E, h-E/M, M) coexist for some range of SNAIL levels, implying that phenotypic heterogeneity and plasticity can occur without active degradation of Zeb-mRNA. On the other hand, in miR200b,c/Zeb circuit, even though the three state coexist for a very narrow range of SNAIL, E to M as well as M to E transition still happens without transitioning to intermediate state, i.e., the **h-E/M state**. A similar behavior persists in the least cooperative versions of miR200a/Zeb (**Fig. 4C**) and miR200b,c/Zeb (**Fig. 4D**) circuits. Further, gradual reduction in translation inhibition cooperativity from six to three decreases h-E/M region significantly in both the least (3; 3,2,2,2) as well as maximal (3;2,1,2,1) cooperative miR200b,c/Zeb circuits compared to miR200a/Zeb least (6;2,1,2,1) and maximal (6;3,2,2,2) cooperative circuits **(Fig. S2 A, B)**. Our results thus suggest that while the absence of active degradation may not prevent the emergence of h-E/M phenotype in miR200a/Zeb circuit and its least cooperative counterpart, it is likely to curtail h-E/M phenotype in miR200b,c/Zeb circuit and its least cooperative counterpart.

**Fig. 4.**
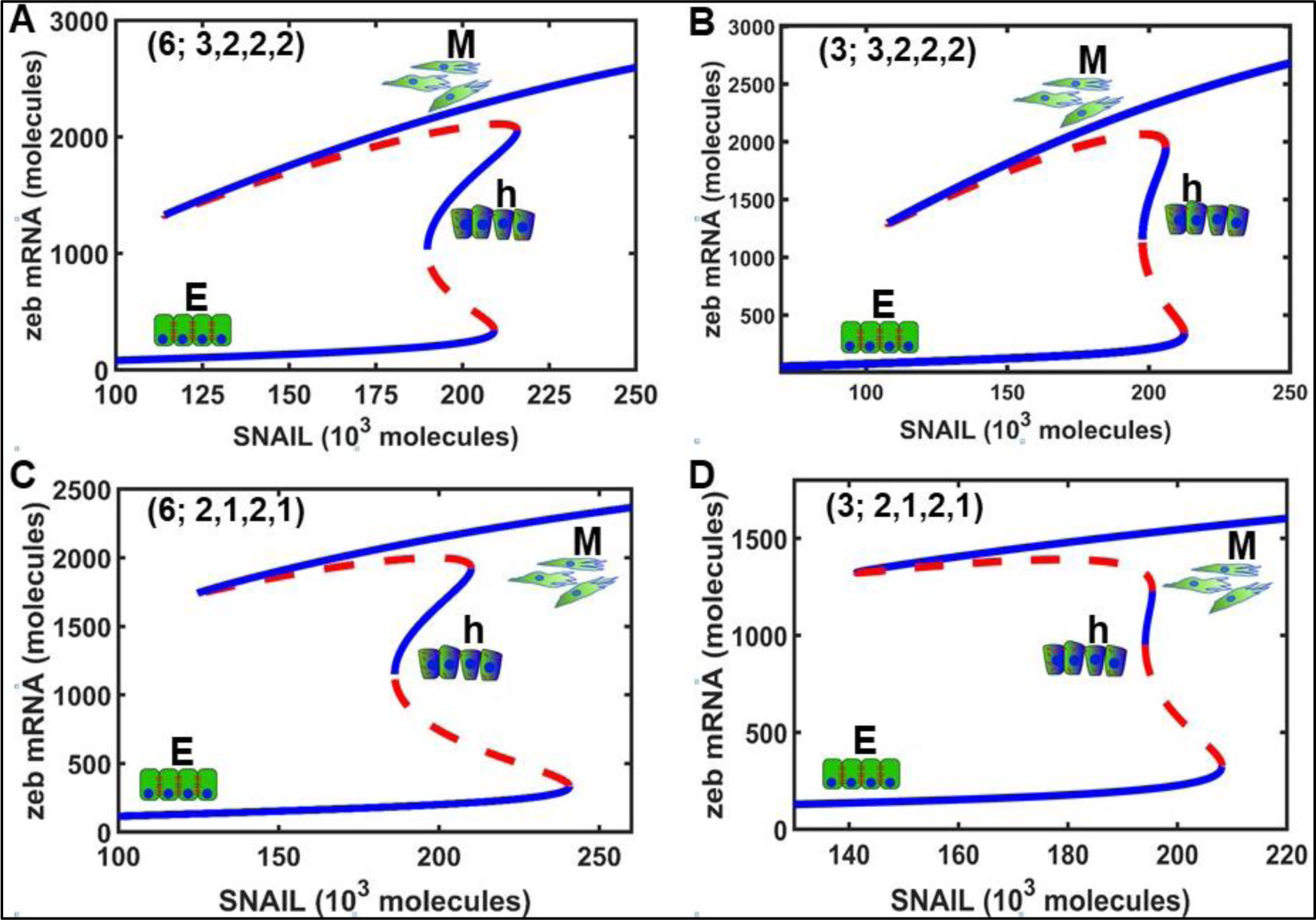
Bifurcation plots showing EMT in the absence of active degradation. (A) EMT in miR200a/Zeb circuit with maximal CC, (B) EMT in miR200b,c/Zeb circuit with maximal CC. (C) and (D) are the least cooperative counter parts of these circuits. Increasing SNAIL levels gradually switches E cells to h-E/M cells and finally to M cells and conversely. Both EMT and MET transitions are symmetric. Also, as cooperativity decreases, h-E/M almost disappears (D), and with further reduction, the circuit may eventually behave as bistable with just E and M phenotypes. (D) Shows that the minimal cooperative circuit to support h-E/M phenotype must have at least three binding sites on Zeb occupied by miR200 (*n* = 3), two binding sites on miR200 occupied by Zeb (*n*_1_ = 2), one binding site each on miR200 and Zeb occupied by SNAIL (*n*_2_ = 1, *n*_4_ = 1), and two binding sites on Zeb occupied by ZEB (*n*_3_ = 2).

### Transcriptional feedback of ZEB can drive EMT in the absence of SNAIL

Many *in vitro* as well as computational studies have reported SNAIL as a key transcriptional regulator driving EMT and metastasis formation. However, a recent *in vivo* study on mouse model of pancreatic ductal adenocarcinomas (PDAC) provided a counter perspective by showing that SNAIL deletion does not block EMT and is thus not rate-limiting for EMT and metastasis^51^. Also, different member of SNAIL family of transcription factors have diverse functionality and not all of them induce EMT. For instance, a recent study reported SNAIL3 as a poor inducer of EMT comforting the concept that the transcription factor functionally diverges from its other family members, SNAIL1 and SNAIL2^52^. This prompted us to investigate whether SNAIL-knocked-out miR200/ZEB circuits, mimicked by model (3), can display EMT. And, if so, what could be the driver of EMT in the absence of SNAIL. Interestingly, our results show that SNAIL is dispensable for EMT and that miR200/Zeb circuit alone can generate and regulate transition of E to M phenotype through intermediate h-E/M phenotype. Specifically, we identify three stable steady states in highly cooperative miR200a/Zeb and miR200b,c/Zeb circuits (**Fig. 5A, B**), as well as in their least cooperative counter parts (**Fig. 5C, D**), while also showing that cooperative self-activation of Zeb (mimicked by *n*_3_ in model (3)) is necessary for tristability.

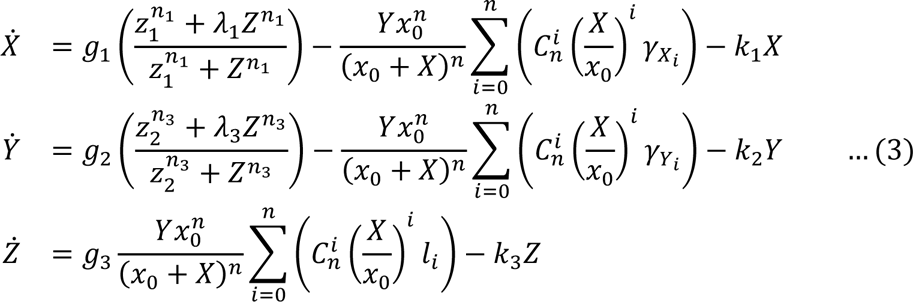

**Fig. 5.**
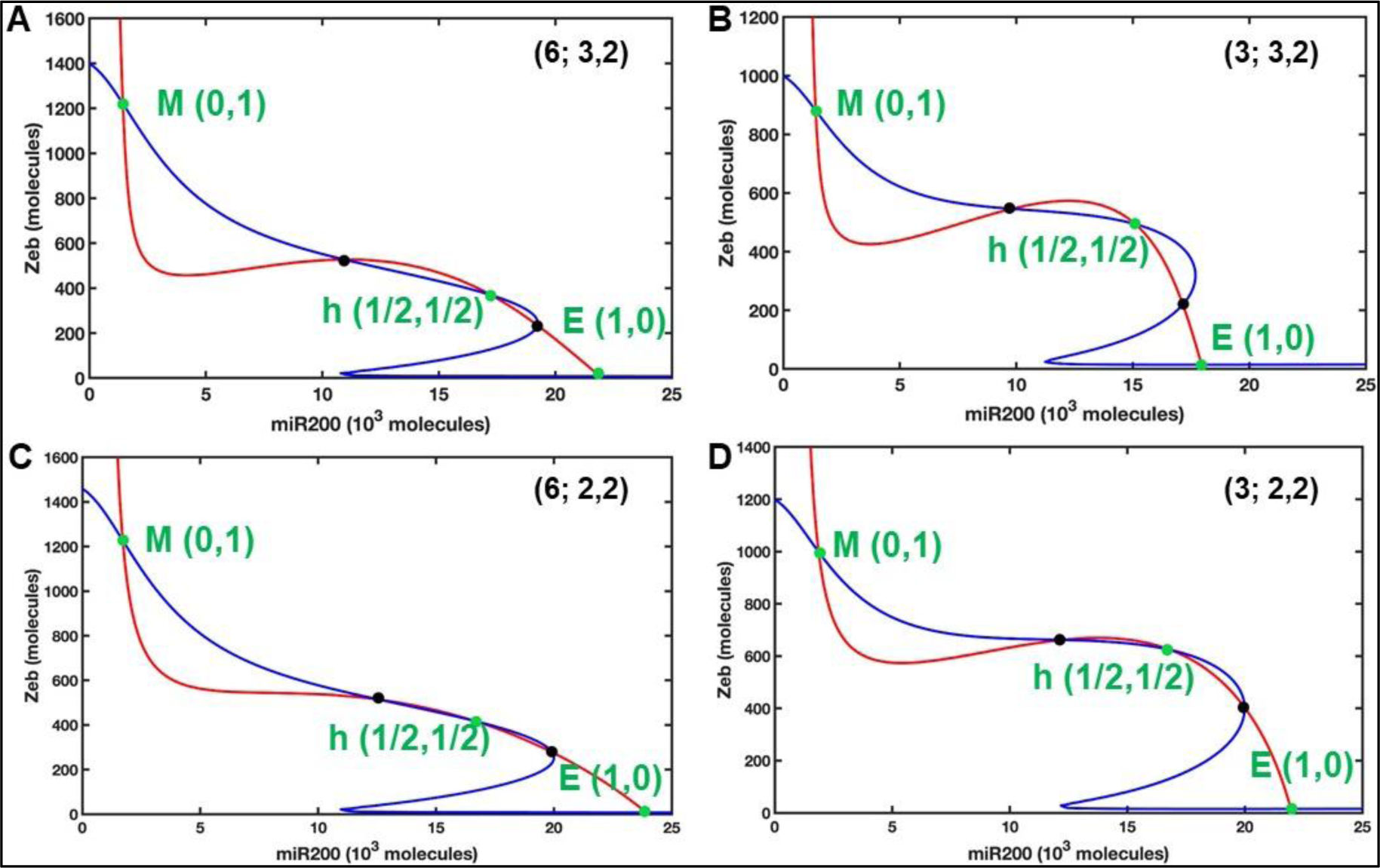
Nullcline plots showing tristability in SNAIL mutated miR200/Zeb circuits. (A) Existence of E, h-E/M, and M phenotypes in miR200a/Zeb circuit with maximal CC: six miR200 binding on Zeb occupied (*n* = 6), three ZEB binding sites on miR200 occupied (*n*_1_ = 3), and two ZEB binding sites on Zeb occupied (*n*_3_ = 2). (B) miR200b,c/Zeb circuit with maximal CC: three miR200 binding on Zeb occupied (*n* = 3), three ZEB binding sites on miR200 occupied (*n*_1_ = 3), and two ZEB binding sites on Zeb occupied (*n*_3_ = 2). (C) and (D) are the least cooperative versions of these circuits. In least cooperative miR200a/Zeb circuit (C), the states are not well-characterized, probably due to mismatch in the number of occupied binding sites at the translation level and transcription level.

To identify the driver of EMT in these circuits, we further demonstrate that in the absence of SNAIL the transcriptional feedback on Zeb (mimicked by *λ*_3_) plays critical role in driving EMT in both the highly cooperative circuits (**Fig. 6A, B**) as well as in the least-cooperative counterparts (**Fig. 6B, C**). However, notably, we found that while the induction of Zeb transcriptional feedback leads to a direct forward transition from E to M phenotype, decreasing the feedback effect gradually leads to backward transition from M to E phenotype through intermediate h-E/M phenotype. This asymmetry in forward and backward transition is partially consistent with the previous experiments showing that active inhibition of SNAIL (mimicked by knockout, in our case) is necessary to initiate direct MET^53^. We further analyzed the effects of variable degradation cooperativities of Zeb due to miR200 by changing n from three to six. We found that such a change does not have any considerable effects on the stability of the three phenotypes in either circuits (Fig. S3 A, B). Our results, thus, identify the redundant role of SNAIL in enabling EMT and uphold the previously recognized role of miR200/ZEB circuit as a “motor of cellular plasticity”.

**Fig. 6.**
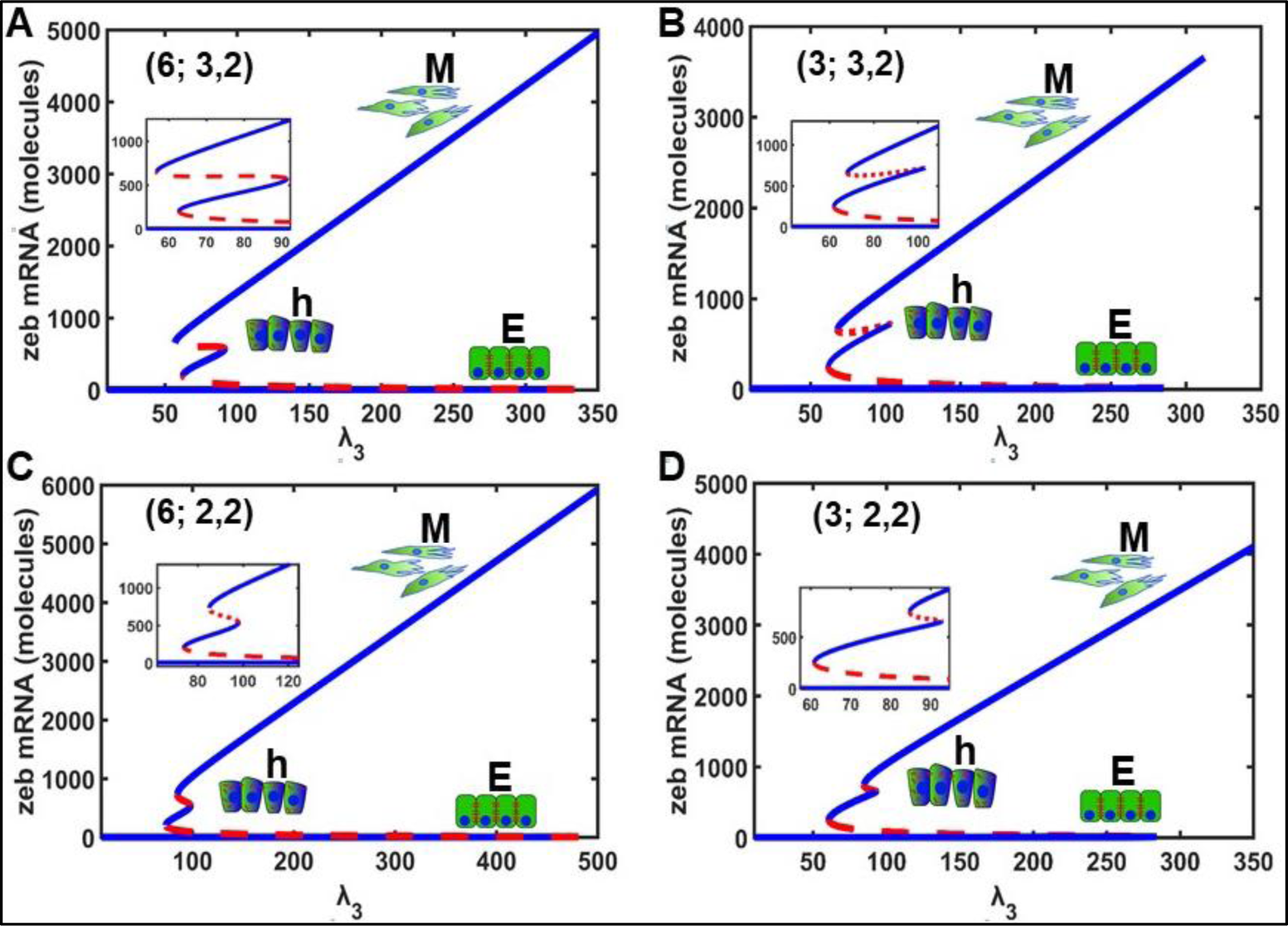
EMT in transcriptional feedback induced miR200/Zeb circuits in the absence of SNAIL. Bifurcation plots showing transcriptional feedback (*λ*_3_) induced phenotypic transition from E to h-E/M to M in miR200a/Zeb (A) and miR200b,c/Zeb (B) circuits with peak CC and in their least cooperative versions (C, D), respectively. Low, intermediate, and high levels of transcriptional feedback on Zeb correspond to E, h-E/M, and M states, respectively. The EMT and MET transition in all the plots, except (A), is symmetric.

### Both SNAIL and Zeb active degradation are dispensable for EMT and tristability

In previous sections, we established that induction of SNAIL and active degradations due to miR200 are separately dispensable for EMT and tristability. To further investigate whether their roles are simultaneously redundant for EMT and tristability, we deleted both SNAIL and active degradation roles in the circuit and obtained a miR200-Zeb self-activated toggle switch whose dynamics can be described by mathematical model in (4). This circuit however differs from the “classic toggle switch” in that one of the interactions occurs at transcription level and the other at translation level. To probe multistable dynamics in this circuit, we made slight changes to two transcriptional parameters (please see Table S1) and identified three stable steady states in both miR200a/Zeb circuit (**Fig. 7A**) as well as in miR200a/Zeb circuit (**Fig. 7B**). Like earlier, based on the expressions levels of SNAIL and Zeb, these states can be clearly categorized into E, h, and M states. Next, to probe the least cooperative (weakly non-linear) versions of either circuits that can produce tri-stability, we found that (*n*_1_=2, *n*_3_=2) combination is the least possible CC in miR200a/Zeb (**Fig. 7C**) that gives tristable dynamics; further decrease leads to bistable dynamics. However, in miR200a/Zeb circuit, the CC can’t be decreased further as we could only have bistability in (3, 2, 2) case. To regain tristability in (*n*_1_=2, *n*_3_=2) case, a translational cooperativity of greater or equal to four is at least required (**Fig. 7D**). Further, it is clear that while tristability is best characterized in both higher and least order cooperative versions of miR200a/Zeb circuits (**Fig. 7A, C**), the same isn’t the case with miR200b,c/Zeb circuit where states appear to be sensitive and can easily switch to bistability (**Fig. 7B, D**). Our results thus show that while simultaneous operation of SNAIL and miR200 assisted active degradation may not be required to attain tristability in miR200a/Zeb circuit, at least one these must be present in miR200b,c/Zeb circuit to achieve tristability. Further, our analysis shows that just three (not six) miR200 binding site on Zeb need to be occupied to achieve tristability and the minimum transcriptional and posttranslational CC to attain tristability is (4; 2, 2), resp.

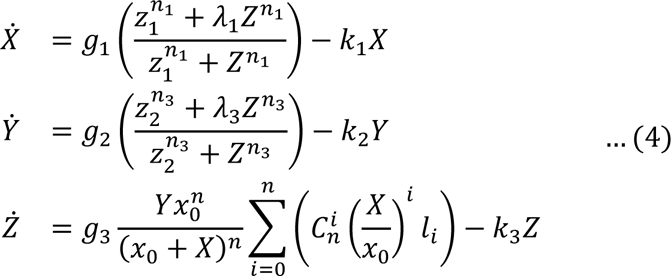

**Fig. 7.**
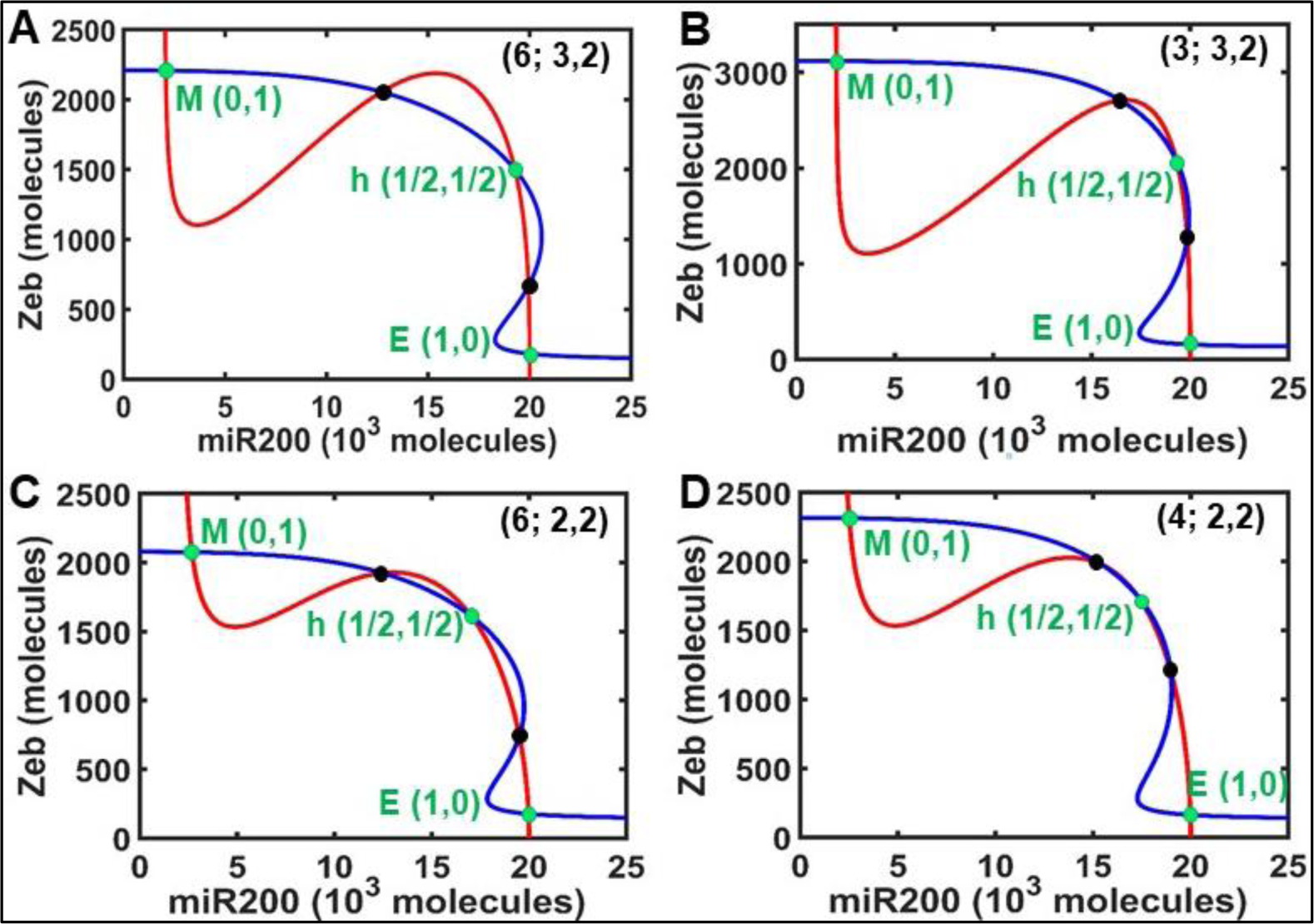
Tristability in SNAIL and active degradation mutated circuits. Nullcline plots showing three phenotypes in highly CC miR200a/Zeb (A) and miR200b,c/Zeb (B) circuits and their least cooperative counterparts (C, D). In the absence of SNAIL and active degradation, at least three miR200 binding sites on Zeb (*n* = 3), three Zeb binding sites on miR200 (*n*_1_ = 3), and two ZEB binding sites on ZEB (*n*_3_ = 2) need to be occupied to exhibit three phenotypes (B). Reduction in cooperativity of Zeb binding to miR200 from *n*_1_ = 3 to *n*_1_ = 2 leads to disappearance of h-E/M state, that is rescued by increasing cooperativity of miR200 binding to Zeb from *n* = 3 to *n* = 4 (D) and is better characterized with further increase to *n* = 6 (C).

Further, to analyze EMT in the absence of SNAIL and miR200 assisted active degradation, we again observe that as the strength of self-activation (*λ*_3_) is increased, both E, h, and M cells emerge in either circuits as shown in **Fig. 8**. However, surprisingly, we found that as *λ*_3_increases, E cells switch directly to M cells without attaining the h-E/M state and as *λ*_3_ decreases, M cells switch directly to E cells. While we also observed direct EMT and MET transition in models without SNAIL or without active degradation, these results contrast with the model in which both are present. In all these models while M to E transition happens directly that any way supports experimental data, E to M transition happens directly only when SNAIL and/or miR200 assisted active degradation is mutated. The same behavior persists upon varying translational cooperativity from three to six in either circuits (**Fig. S4, A, B**). Our results thus link occurrence of intermediate h-E/M phenotype during EMT with SNAIL induction and/or miR200 assisted active degradation and offer crucial insights to curtail intermediate phenotype during EMT and prevent heterogeneity.

**Fig. 8.**
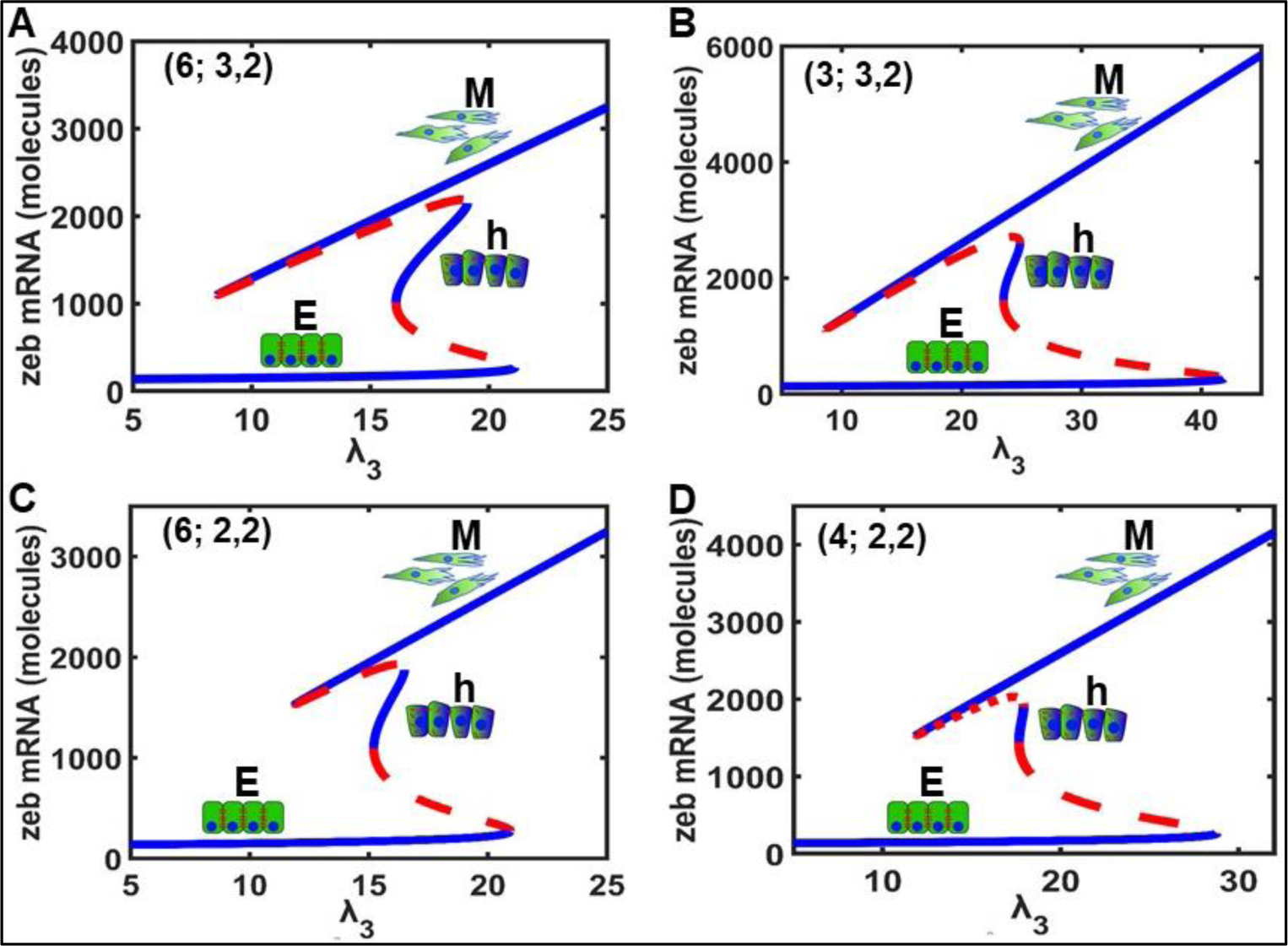
EMT in SNAIL and active degradation mutated circuits. Bifurcation plots show cell state transition from E to h-E/M to M state upon induction of Zeb transcriptional feedback in highly cooperative miR200a/Zeb (A) and miR200b,c/Zeb circuits and their least cooperative counterparts (C,D), resp. In (D) h-E/M state occurs for narrow input range, showing that CC like (*n*; *n*_1_, *n*_3_) = (3; 2, 2) is insufficient to support h-E/M phenotype (D). Also, (A) shows the best characterization of the three states, implying that higher order cooperative binding can support tristability in the absence of SNAIL and active degradation.

## Discussion and Conclusion

Cell fate switching enabled by EMT drives carcinoma cell phenotypic plasticity by conferring certain mesenchymal traits on epithelial cells. EMT activated cells often proceed only partway towards a fully mesenchymal state and are arrayed at various points along the epithelial (E) to mesenchymal (M) phenotypic axis, leading to non-genetic phenotypic heterogeneity. Phenotypes that emerge due to partial activation of EMT are reported to be critical for metastasis, aggression of cancers, and therapy failure. Therefore identifying inhibitors of partial EMT is crucial to the success of future therapies aiming to curtail metastasis.

Gene networks underlying EMT of widely different sizes have been employed to usefully model the regulation of choice between E and M fates. However, the most extensively studied among these networks is the miR200/Zeb mutually inhibitory feedback loop. Numerous experimental and computational studies point to the miR200/Zeb1 axis as crucial in regulating binary choice between E and M phenotypes and together with SNAIL, the circuit is referred as a molecular motor of cellular plasticity in metastasis. Apparently, miR200/Zeb circuit is not complex as it involves just two molecules with few binding interactions. However, combinatorial cooperativity makes this circuit inherently complex and highly nonlinear. Further, the presence of miR200 in this circuit adds to mechanistic complexity at the translation level, since microRNA’s can silence mRNA’s by translation inhibition as well as by active degradation. Thus unravelling the implications of CC as well as miR200 silencing effects in phenotypic plasticity is complex to determine. Further, Zeb has different binding cooperativities for different families of miR200: six for miR200a and three each for miR200b and miR200c molecules. Whether CC has any role in phenotypic heterogeneity and in EMT regulation remains uninvestigated until now. Previous studies have exploited maximum CC to demonstrate that SNAIL induced miR200a/Zeb circuit acts as a ternary switch, allowing intermediate h E/M phenotype, apart from E and M phenotypes. A natural question arises here: does every binding site on each gene need to be occupied to attain three phenotypes; if not, what is the minimum number for each gene? Also, are there any phenotypic consequences corresponding to different binding events?

To answer these questions, we analysed steady state dynamics of miR200a/Zeb and miR200a/Zeb circuits under different cooperative conditions. Our results demonstrate that h-E/M phenotype can occur without higher order cooperative binding of molecules at the transcription and translation level in both the circuits. We found a minimum threshold of binding cooperativity at both the transcription and translation levels that must be fulfilled to achieve tristability: Zeb to miR200 binding and Zeb self-activation binding must have cooperativity of order at least two and miR200 binding to Zeb must have cooperativity of order at least three. Cooperativities below these thresholds do not support h-E/M state and thus circuits give bistable dynamics. To understand the connection of miR200 silencing effects with tristability, we decoupled the simultaneous translation inhibition and active degradation roles of Zeb due to miR200. Unlike previous studies, we found that while translation inhibition is necessary for the hybrid phenotype, active degradation seems to be dispensable. Further, we show that the role of SNAIL in the phenotypic transition in EMT can be redundant and in the absence of SNAIL, the strength of cooperative transcriptional feedback on Zeb can control EMT: increasing Zeb feedback strength drives cells from epithelial to hybrid and finally to mesenchymal states. Our findings are in line with the experimental data reporting induction of EMT without SNAIL^10^ and mutating miR200 binding sites on Zeb to induce EMT^39^. Our results thus untangle the molecular basis of EMT regulation and uphold the SNAIL-independent role of miR200/Zeb circuit as a crucial network in controlling cellular plasticity between E, h-E/M and M phenotypes through the (in)activation of its different mechanism of operation. These results may provide unconventional alternatives to prevent phenotypic transition due to EMT in order to curtail the progression of carcinomas.

## Author Contributions

**Mubasher Rashid**: Conceptualization, methodology, data analysis, writing, supervision, funding acquisition.

**Brasanna M Devi and Malay Banerjee**: Data analysis.

## Methods

**Steady state analysis**: Maple (release 2023.0) was used to identify feasible steady states of all models in the study. The stability of each steady state was verified using Jacobian method.

**Bifurcation analysis:** All bifurcation were performed using MATCONT (version 7.1) as GUI in MATLAB (version 9.9.0.1857802 (R2020b)). For each model, bifurcations (both forward and backward) were initiated around the equilibrium points calculated in MAPLE.

## Acknowledgements

We benefited from useful discussions with Prof. Mohit Kumar Jolly, Center for BioSystems Science and Engineering, Indian Institute of Science, Bangalore, India. We gratefully acknowledge funding by the Department of Science and Technology (DST), Govt. of India, to MR (grant number: DST/INSPIRE/04/2020/001492).

## Supplementary material

Table S1 lists the parmeter values for models (1) - (4). Since the mechanims associated to these parameters operate at transcription level of biological regulation, we call them transcriptional kinetic parameters. TI denotes translation inhibition and AD denotes active degradation. *“NA”* denotes non-involvement of the parameter in the particular model. “K” denotes thousand.

**Table S1:** Transcriptional kinetic parameters for the four models.

| **Table S1: Transcriptional kinetic parameters for the four models.** |  |  |  |  |
| --- | --- | --- | --- | --- |
| **Parameter** | **Models and corresponding parameter values** |  |  |  |
|  | **Model (1):**<br>**Both TI and**<br>**AD, with**<br>**SNAIL** | **Model (2):**<br>**TI only, with**<br>**SNAIL** | **Model (3):**<br>**Both TI and**<br>**AD, without**<br>**SNAIL** | **Model (4):**<br>**TI only, without**<br>**SNAIL** |
| $g_1$ (conc. $h^{-1}$) | 1500 | 1500 | 1500 | 1000 |
| $g_2$ (conc. $h^{-1}$) | 12.5 | 12.5 | 15 | 65 |
| $g_3$ ($h^{-1}$) | 100 | 100 | 100 | 100 |
| $z_1$ (conc.) | 200K | 200K | 200K | 200K |
| $z_2$ (conc.) | 24K | 50K | 25K | 50K |
| $s_1$ (conc.) | 180K | 180K | NA | NA |
| $s_2$ (conc.) | 180K | 180K | NA | NA |
| $\lambda_1$ | 0.1 | 0.1 | 0.1 | 0.1 |
| $\lambda_2$ | 0.8 | 0.4 | NA | NA |
| $\lambda_3$ | 12.5 | 15 | 70 | 15 |
| $\lambda_4$ | 9.0 | 10 | NA | NA |
| $k_1$ ($h^{-1}$) | 0.05 | 0.05 | 0.05 | 0.05 |
| $k_2$ ($h^{-1}$) | 0.5 | 0.5 | 0.5 | 0.5 |
| $k_3$ ($h^{-1}$) | 0.1 | 0.1 | 0.1 | 0.1 |
| $x_0$ | 10K | 10K | 10K | 10K |
| S | 200K | 200K | NA | NA |

Table S2 lists the posttranscriptional parameters involved in models (1) – (4). Here, 0 – 6 denote the number of miR200 molecules bound to Zeb. With each occupied miR200 binding site, a degradation rate is assigned to miR200 (*γ_X_i__*) and Zeb (*γ_Y_i__*) in the corresponding miR200-Zeb complex. The translation rate of Zeb (to form ZEB) is modelled with *l*_*i*_. It may be noted that degradation as well as translation rates are assumed to saturare after Zeb binds three or more molecules of miR200. As shown in the table, models (2) and (4) don’t contain degradation rates for miR200 and Zeb, because these models are studied without the role of miR200 active degradation.

**Table S2:** Post-transcription kinetic parameters.

| **Table S2: Post-transcription kinetic parameters** |  |  |  |  |  |  |  |  |
| --- | --- | --- | --- | --- | --- | --- | --- | --- |
| **Model Name** |  | **No. of miR200 binding sites on Zeb, n (i = 0, 1, ..., 6)** |  |  |  |  |  |  |
|  |  | 0 | 1 | 2 | 3 | 4 | 5 | 6 |
| **Model (1)** | $\gamma_{x_i} (h^{-1})$ | 0 | 0.007 | 0.07 | 0.7 | 0.7 | 0.7 | 0.7 |
| | $\gamma_{y_i} (h^{-1})$ | 0 | 0.04 | 0.2 | 1 | 1 | 1 | 1 |
| | $l_i$ | 1 | 0.6 | 0.3 | 0.1 | 0.05 | 0.05 | 0.05 |
| **Model (2)** | $l_i$ | 1 | 0.5 | 0.2 | 0.02 | 0.02 | 0.02 | 0.02 |
| **Model (3)** | $\gamma_{x_i} (h^{-1})$ | 0 | 0.005 | 0.05 | 0.5 | 0.5 | 0.5 | 0.5 |
| | $\gamma_{y_i} (h^{-1})$ | 0 | 0.04 | 0.2 | 1 | 1 | 1 | 1 |
| | $l_i$ | 1 | 0.6 | 0.3 | 0.1 | 0.05 | 0.05 | 0.05 |
| **Model (4)** | $l_i$ | 1 | 0.5 | 0.2 | 0.02 | 0.02 | 0.02 | 0.02 |

### Tristability in model (2): miR200a/Zeb and miR200b,c/Zeb networks without active degradation

**Fig. S1.**
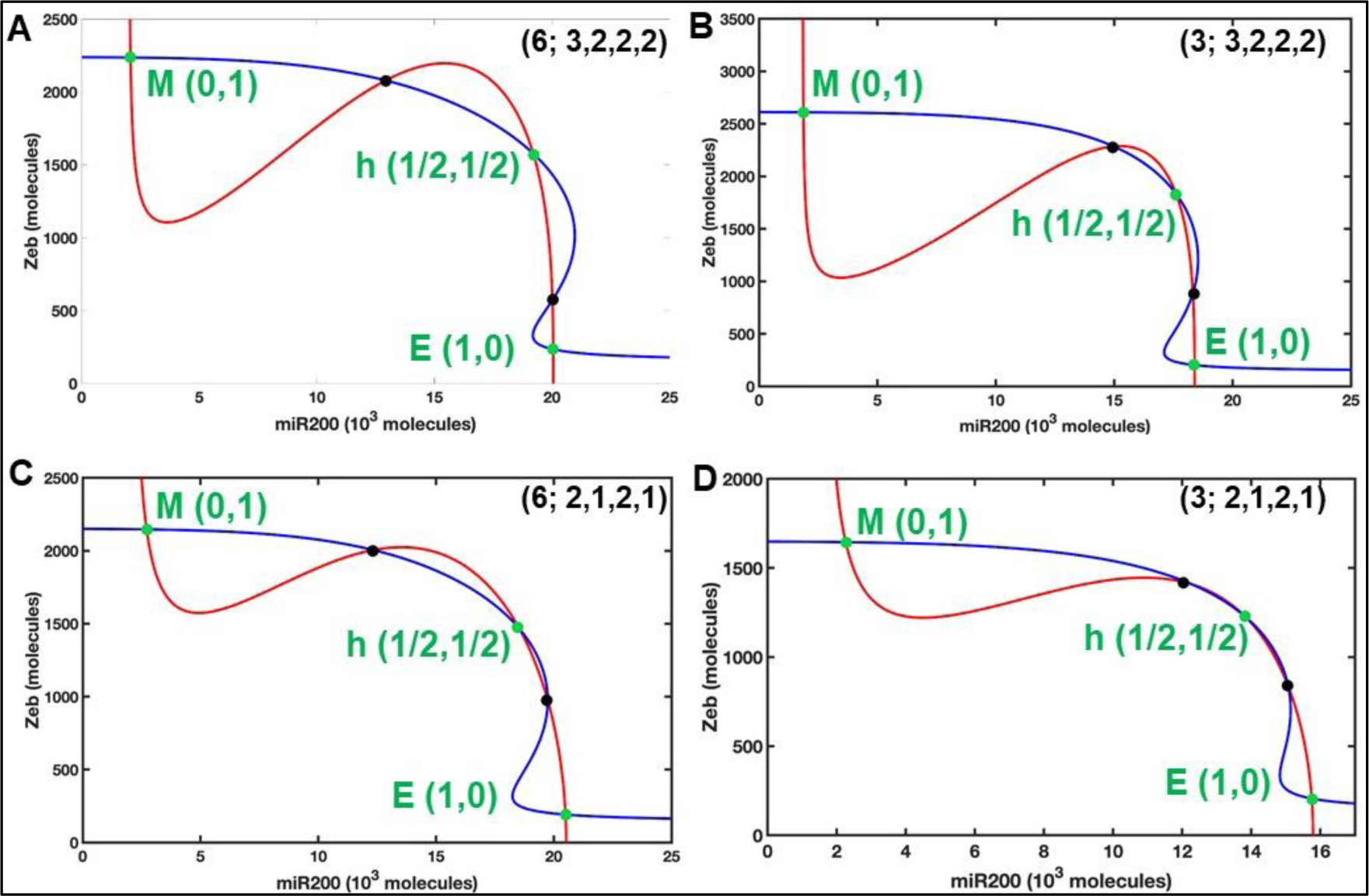
Nullcline plots of miR200/Zeb circuits without active degradation. Tristability in miR200a/Zeb (A) and miR200b,c/Zeb (B) circuits with all binding sites occupied (i.e. maximal CC) and that in their least cooperative counterparts (C, D), resp. The five intersections represent five steady states out of which three (green dots) are stable and two (black dots) are unstable. High (low) values of miR200 (Zeb) corresponds to E state, intermediate values of miR200 (Zeb) correspond to hybrid state and low (high) values of miR200 (Zeb) corresponds to M state.

### EMT in model (2) under different translational cooperativity conditions

EMT in SNAIL induced miR200a/Zeb and miR200b,c/Zeb circuits without considering the role of active degradations. (A) as the translational cooperativity is varied between three to six, we gradually move from highly cooperative miR200b,c/Zeb (3; 3,2,2,2) circuit to highly cooperative miR200a/Zeb (6;3,2,2,2) circuit. Similarly, in (B), we move from least cooperative miR200b,c/Zeb (3; 2,1,2,1) circuit to least cooperative miR200a/Zeb (6; 2,1,2,1) circuit. In either case, it is evident that hybrid state (phenotype) appears to be more stable and emerges for wider levels of SNAIL values in the case of maximal translational cooperative (6, red curve) circuits, compared to their least cooperative counterparts (3, blue). This shows that in the absence of active degradation, a highly cooperativity translational inhibition is required to compensate the lost affects of active degradation.

**Fig. S2.**
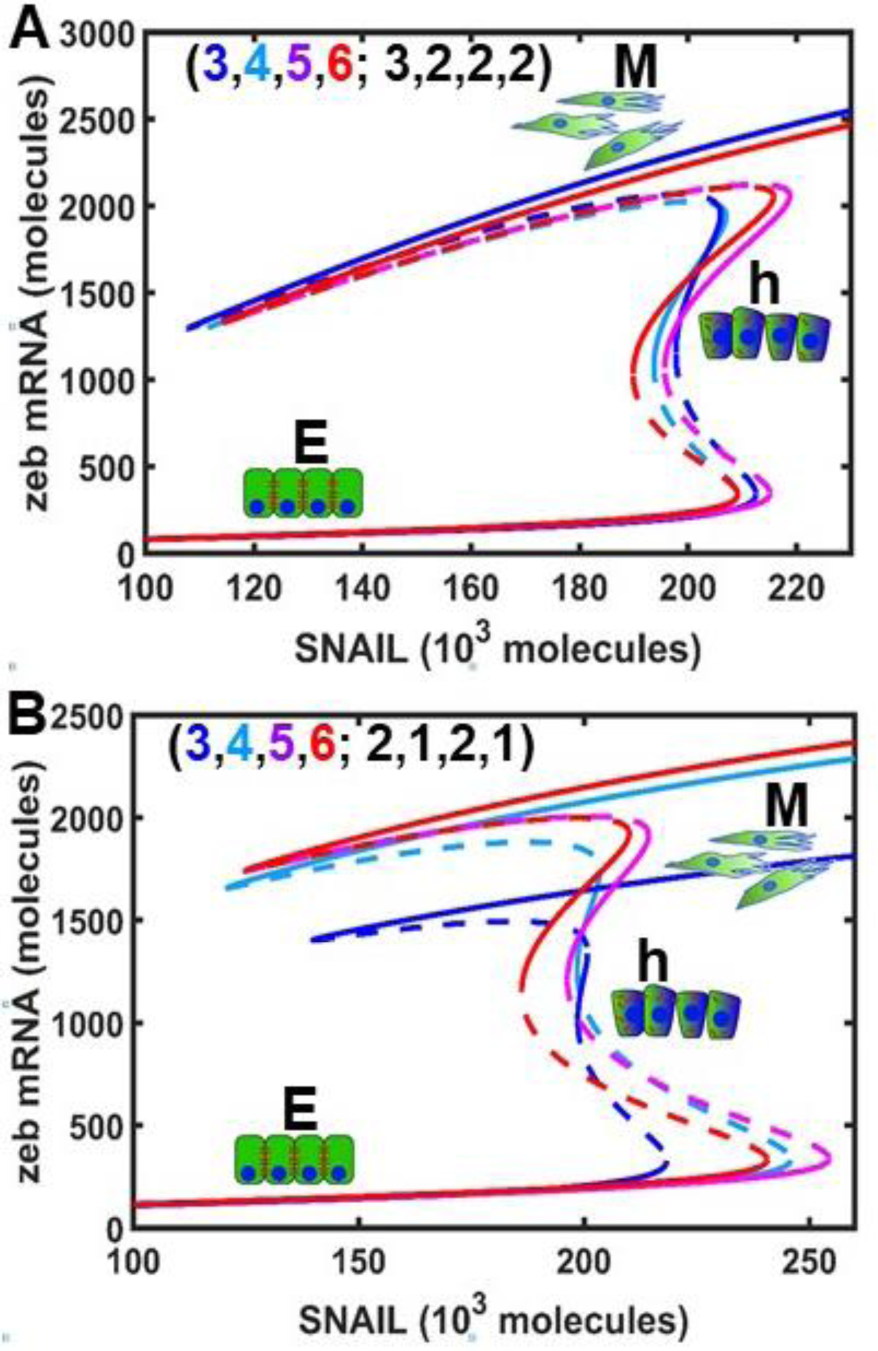
Bifurcation plots showing EMT for different translational cooperativity conditions. (A) Transcriptional cooperativity is fixed at its maximum value (3,2,2,2) and posttranscriptional (or translational) cooperativity of miR200 binding to Zeb is varied between three to six. (B) same as previous case, but here the transcriptional cooperativity is fixed at minimal value (2,1,2,1). Solid curves represent stable states and dotted curves represent unstable states

### EMT in model (3) under different translational cooperativity conditions

EMT in SNAIL knock-out least maximum (Fig. S3 A, red) and minimum (Fig. S3 B, red) cooperative miR200a/Zeb circuit and that in maximumum (Fig. S3 A, blue) and minimum (Fig. S3 B, blue) cooperative miR200b,c/Zeb circuits. The plots show that as the strength of transcriptional feedback increases, E cells switch to h-E/M cells and a continuous induction further switches h-E/M cells to M cells to achieve a complete EMT. Also, As the transcriptional feedback decreases, MET through h-E/M state is observed. It may be noted that both EMT and MET are symmetric, meaning that both procees via h-E/M states. Also, in both maximal and least cooperative circuits, variable translational cooperativity of Zeb active degradation due to miR200 has no affect on the stability of h-E/M phenotype, since the h-E/M phenotype is equally stable for all values between three to six. Since we observed in model (2) that active degradation deletion, affects the stability of h-E/M phenotype and maximal cooperativity of translational inhibition was able to regain its stability at that time, we conclude that active degradation of Zeb due to miR200 might be one of the important deciding factors for the emergence of h-E/M phenotype in either circuits.

**Fig. S3.**
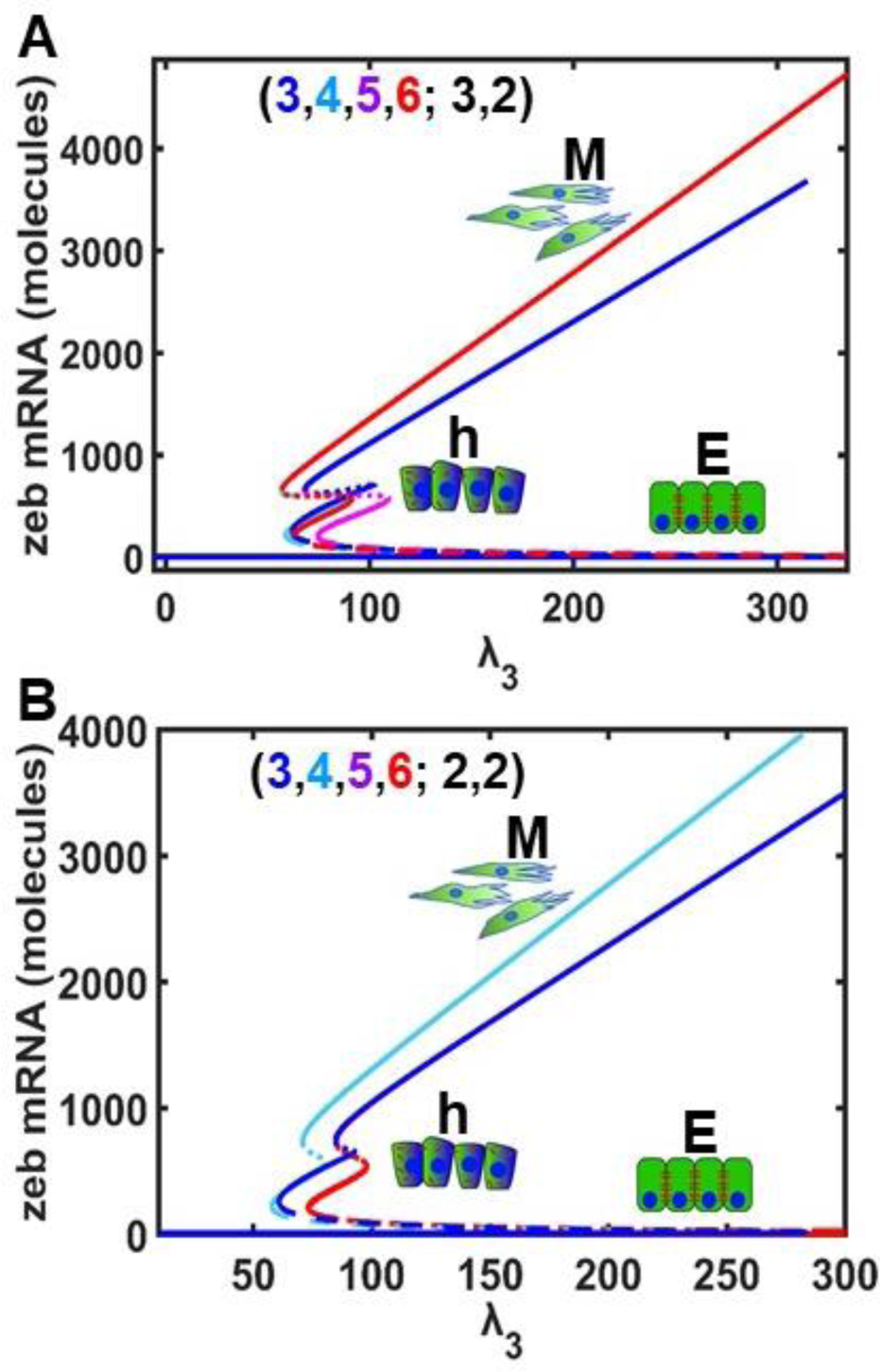
Bifurcation plots showing EMT induced by transcriptional feedback (*λ*_3_) on Zeb. (A) transcriptional cooperativity is fixed at maximal value (3,2) and translational cooperativity is varied between three to six, so that (3; 3,2) and (6;3,2) correspond to highly cooperative miR200b,c/Zeb and miR200a/Zeb circuits, resp. (B) Same as previous case, except that the circuits are least cooperative.

### EMT in model (4) under different translational cooperativity conditions

In this model both active degrations due to miR200 and the affects of SNAIL are knocked out. Varying translational cooperativity (cooperativity of translation inhibition of Zeb due to miR200) doesn’t have significant affects on the stability of h-E/M phenotype. As cooperative feedback strength (*λ*_3_) is increased/decreased EMT/MET is observed. It is evident from the plots that both EMT and MET are symmetric, meaning that the switch happens directly from E to M (in the forward case) and M to E (in the reverse case) phenotypes without transitioning to the intermediate h-E/M state. Fig S4 (B) shows that ZEB binding to miR200 and ZEB binding to Zeb must have a cooperativity of order at least two (n_1_=n_3_=2) to attain tristability. Thus the occurrence of h-E/M phenotype depends on the cooperative transcriptional feedback activation of Zeb as well as the transcriptional inhibition of miR200 by ZEB transcription factor.

**Fig. S4.**
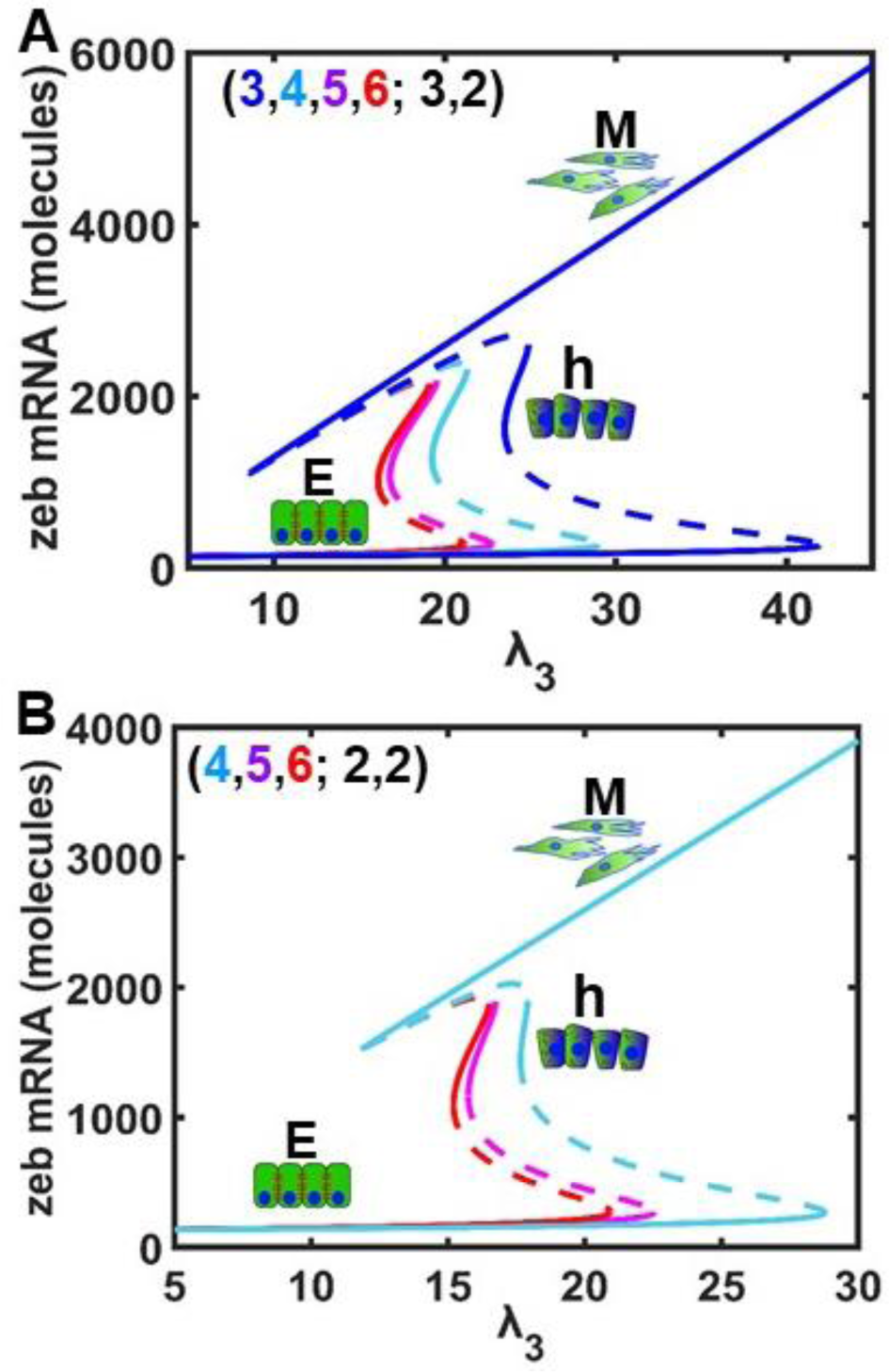
Bifurcation plots showing EMT induced by transcriptional feedback (*λ*_3_) on Zeb. (A) transcriptional cooperativity is fixed at maximal value (3,2) and translational cooperativity is varied between three to six, so that (3; 3,2) and (6;3,2) correspond to highly cooperative miR200b,c/Zeb and miR200a/Zeb circuits, resp. (B) Same as previous case, except that the transcriptional cooperativity is fixed at minimum value (2,2).

## References

1. Fares J, Fares MY, Khachfe HH, Salhab HA, Fares Y. Molecular principles of metastasis: a hallmark of cancer revisited. Signal Transduct Target Ther. 2020;5(1). doi:10.1038/s41392-020-0134-x

2. Fuqua SAW. Metastasis: Complexity thwarts precision targeting. British Journal of Cancer 2021 125:8. 2021;125(8):1033-1035. doi:10.1038/s41416-021-01401-1

3. Ye X, Weinberg RA. Epithelial–Mesenchymal Plasticity: A Central Regulator of Cancer Progression. Trends Cell Biol. 2015;25(11):675–686. doi:10.1016/J.TCB.2015.07.012

4. Thiery JP, Acloque H, Huang RYJ, Nieto MA. Epithelial-Mesenchymal Transitions in Development and Disease. Cell. 2009;139(5):871–890. doi:10.1016/J.CELL.2009.11.007

5. Bastid J. EMT in carcinoma progression and dissemination: Facts, unanswered questions, and clinical considerations. Cancer and Metastasis Reviews. 2012;31(1-2):277–283. doi:10.1007/S10555-011-9344-6/FIGURES/2

6. Aiello NM, Kang Y. Context-dependent EMT programs in cancer metastasis. Journal of Experimental Medicine. 2019;216(5):1016–1026. doi:10.1084/JEM.20181827

7. Lambert AW, Pattabiraman DR, Weinberg RA. Emerging Biological Principles of Metastasis. Cell. 2017;168(4):670–691. doi:10.1016/J.CELL.2016.11.037

8. Mani SA, Guo W, Liao MJ, et al. The Epithelial-Mesenchymal Transition Generates Cells with Properties of Stem Cells. Cell. 2008;133(4):704–715. doi:10.1016/J.CELL.2008.03.027

9. Lambert AW, Weinberg RA. Linking EMT programmes to normal and neoplastic epithelial stem cells. Nature Reviews Cancer 2021 21:5. 2021;21(5):325-338. doi:10.1038/s41568-021-00332-6

10. Zheng X, Carstens JL, Kim J, et al. Epithelial-to-mesenchymal transition is dispensable for metastasis but induces chemoresistance in pancreatic cancer. Nature. 2015;527(7579):525-530. doi:10.1038/nature16064

11. Singh A, Settleman J. EMT, cancer stem cells and drug resistance: an emerging axis of evil in the war on cancer. Oncogene 2010 29:34. 2010;29(34):4741-4751. doi:10.1038/onc.2010.215

12. Saxena M, Stephens MA, Pathak H, Rangarajan A. Transcription factors that mediate epithelial–mesenchymal transition lead to multidrug resistance by upregulating ABC transporters. Cell Death & Disease 2011 2:7. 2011;2(7):e179-e179. doi:10.1038/cddis.2011.61

13. Vega S, Morales A V., Ocaña OH, Valdés F, Fabregat I, Nieto MA. Snail blocks the cell cycle and confers resistance to cell death. Genes Dev. 2004;18(10):1131–1143. doi:10.1101/GAD.294104

14. Lu W, Kang Y. Epithelial-Mesenchymal Plasticity in Cancer Progression and Metastasis. Dev Cell. 2019;49(3):361–374. doi:10.1016/j.devcel.2019.04.010

15. Gupta PB, Pastushenko I, Skibinski A, Blanpain C, Kuperwasser C. Phenotypic Plasticity: Driver of Cancer Initiation, Progression, and Therapy Resistance. Cell Stem Cell. 2019;24(1). doi:10.1016/j.stem.2018.11.011

16. Kröger C, Afeyan A, Mraz J, et al. Acquisition of a hybrid E/M state is essential for tumorigenicity of basal breast cancer cells. Proc Natl Acad Sci U S A. 2019;116(15):7353–7362. doi:10.1073/pnas.1812876116

17. Jolly MK, Mani SA, Levine H. Hybrid epithelial/mesenchymal phenotype(s): The ‘fittest’ for metastasis? Biochim Biophys Acta Rev Cancer. 2018;1870(2):151–157. doi:10.1016/j.bbcan.2018.07.001

18. Pastushenko I, Blanpain C. EMT Transition States during Tumor Progression and Metastasis. Trends Cell Biol. 2019;29(3). doi:10.1016/j.tcb.2018.12.001

19. Brooks MD, Burness ML, Wicha MS. Therapeutic Implications of Cellular Heterogeneity and Plasticity in Breast Cancer. Cell Stem Cell. 2015;17(3):260–271. doi:10.1016/J.STEM.2015.08.014

20. Liao TT, Yang MH. Hybrid Epithelial/Mesenchymal State in Cancer Metastasis: Clinical Significance and Regulatory Mechanisms. Cells. 2020;9(3). doi:10.3390/cells9030623

21. Jolly MK, Somarelli JA, Sheth M, et al. Hybrid epithelial/mesenchymal phenotypes promote metastasis and therapy resistance across carcinomas. Pharmacol Ther. 2019;194. doi:10.1016/j.pharmthera.2018.09.007

22. Jolly MK, Boareto M, Huang B, et al. Implications of the hybrid epithelial/mesenchymal phenotype in metastasis. Front Oncol. 2015;5(JUN). doi:10.3389/fonc.2015.00155

23. Zhang Y, Donaher JL, Das S, et al. Genome-wide CRISPR screen identifies PRC2 and KMT2D-COMPASS as regulators of distinct EMT trajectories that contribute differentially to metastasis. Nat Cell Biol. 2022;24(4):554–564. doi:10.1038/s41556-022-00877-0

24. Rozum J, Albert R. Leveraging network structure in nonlinear control. NPJ Syst Biol Appl. 2022;8(1). doi:10.1038/s41540-022-00249-2

25. Charitou T, Bryan K, Lynn DJ. Using biological networks to integrate, visualize and analyze genomics data. Genetics Selection Evolution. 2016;48(1). doi:10.1186/s12711-016-0205-1

26. Sampson VB, David JM, Puig I, et al. Wilms’ tumor protein induces an epithelial-mesenchymal hybrid differentiation state in clear cell renal cell carcinoma. PLoS One. 2014;9(7). doi:10.1371/journal.pone.0102041

27. Jolly MK, Tripathi SC, Jia D, et al. Stability of the hybrid epithelial/mesenchymal phenotype. Oncotarget. 2016;7(19). doi:10.18632/oncotarget.8166

28. Subbalakshmi AR, Kundnani D, Biswas K, et al. NFATc Acts as a Non-Canonical Phenotypic Stability Factor for a Hybrid Epithelial/Mesenchymal Phenotype. Front Oncol. 2020;10. doi:10.3389/fonc.2020.553342

29. Tian XJ, Zhang H, Xing J. Coupled reversible and irreversible bistable switches underlying TGFβ-induced epithelial to mesenchymal transition. Biophys J. 2013;105(4). doi:10.1016/j.bpj.2013.07.011

30. Tripathi S, Kessler DA, Levine H, Balázsi G, Albert R. Minimal frustration underlies the usefulness of incomplete regulatory network models in biology. Published online 2022. doi:10.1073/pnas

31. Rashid M, Hari K, Thampi J, Santhosh NK, Jolly MK. Network topology metrics explaining enrichment of hybrid epithelial/mesenchymal phenotypes in metastasis. Faeder JR, ed. PLoS Comput Biol. 2022;18(11):e1010687. doi:10.1371/journal.pcbi.1010687

32. Hari K, Sabuwala B, Subramani BV, et al. Identifying inhibitors of epithelial–mesenchymal plasticity using a network topology-based approach. NPJ Syst Biol Appl. 2020;6(1). doi:10.1038/s41540-020-0132-1

33. Korpal M, Kang Y. The emerging role of miR-200 family of microRNAs in epithelial-mesenchymal transition and cancer metastasis. RNA Biol. 2008;5(3). doi:10.4161/rna.5.3.6558

34. Park SM, Gaur AB, Lengyel E, Peter ME. The miR-200 family determines the epithelial phenotype of cancer cells by targeting the E-cadherin repressors ZEB1 and ZEB2. Genes Dev. 2008;22(7). doi:10.1101/gad.1640608

35. Larsen JE, Nathan V, Osborne JK, et al. ZEB1 drives epithelial-to-mesenchymal transition in lung cancer. Journal of Clinical Investigation. 2016;126(9). doi:10.1172/JCI76725

36. Kurahara H, Takao S, Maemura K, et al. Epithelial-mesenchymal transition and mesenchymal-epithelial transition via regulation of ZEB-1 and ZEB-2 expression in pancreatic cancer. J Surg Oncol. 2012;105(7). doi:10.1002/jso.23020

37. Brabletz S, Brabletz T. The ZEB/miR-200 feedback loop-a motor of cellular plasticity in development and cancer? EMBO Rep. 2010;11(9):670–677. doi:10.1038/embor.2010.117

38. Chaffer CL, Marjanovic ND, Lee T, et al. Poised Chromatin at the ZEB1 Promoter Enables Breast Cancer Cell Plasticity and Enhances Tumorigenicity. Cell. 2013;154(1):61–74. doi:10.1016/J.CELL.2013.06.005

39. Title AC, Hong SJ, Pires ND, et al. Genetic dissection of the miR-200–Zeb1 axis reveals its importance in tumor differentiation and invasion. Nat Commun. 2018;9(1). doi:10.1038/s41467-018-07130-z

40. Hill L, Browne G, Tulchinsky E. ZEB/miR-200 feedback loop: At the crossroads of signal transduction in cancer. Int J Cancer. 2012;132(4):745–754. doi:10.1002/ijc.27708

41. Brabletz S, Bajdak K, Meidhof S, et al. The ZEB1/miR-200 feedback loop controls Notch signalling in cancer cells. EMBO Journal. 2011;30(4). doi:10.1038/emboj.2010.349

42. Bu P, Chen KY, Chen JH, et al. A microRNA miR-34a-regulated bimodal switch targets notch in colon cancer stem cells. Cell Stem Cell. 2013;12(5). doi:10.1016/j.stem.2013.03.002

43. Lu M, Jolly MK, Gomoto R, Huang B, Onuchic J, Ben-Jacob E. Tristability in cancer-associated microRNA-TF chimera toggle switch. Journal of Physical Chemistry B. 2013;117(42):13164–13174. doi:10.1021/jp403156m

44. Lu M, Jolly MK, Levine H, Onuchic JN, Ben-Jacob E. MicroRNA-based regulation of epithelial-hybrid-mesenchymal fate determination. Proc Natl Acad Sci U S A. 2013;110(45):18144–18149. doi:10.1073/pnas.1318192110

45. Burk U, Schubert J, Wellner U, et al. A reciprocal repression between ZEB1 and members of the miR-200 family promotes EMT and invasion in cancer cells. EMBO Rep. 2008;9(6). doi:10.1038/embor.2008.74

46. Guaita S, Puig I, Francí C, et al. Snail induction of epithelial to mesenchymal transition in tumor cells is accompanied by MUC1 repression and ZEB1 expression. Journal of Biological Chemistry. 2002;277(42). doi:10.1074/jbc.M206400200

47. Levine E, Jacob E Ben, Levine H. Target-specific and global effectors in gene regulation by microRNA. Biophys J. 2007;93(11). doi:10.1529/biophysj.107.118448

48. Zhang J, Tian XJ, Zhang H, et al. TGF-β-induced epithelial-to-mesenchymal transition proceeds through stepwise activation of multiple feedback loops. Sci Signal. 2014;7(345):ra91. doi:10.1126/SCISIGNAL.2005304/SUPPL_FILE/7_RA91_SM.PDF

49. Samavarchi-Tehrani P, Golipour A, David L, et al. Functional Genomics Reveals a BMP-Driven Mesenchymal-to-Epithelial Transition in the Initiation of Somatic Cell Reprogramming. doi:10.1016/j.stem.2010.04.015

50. Li R, Liang J, Ni S, et al. A Mesenchymal-to-Epithelial Transition Initiates and Is Required for the Nuclear Reprogramming of Mouse Fibroblasts. doi:10.1016/j.stem.2010.04.014

51. Paul MC, Schneeweis C, Falcomatà C, et al. Non-canonical functions of SNAIL drive context-specific cancer progression. Nature Communications 2023 14:1. 2023;14(1):1-21. doi:10.1038/s41467-023-36505-0

52. Gras B, Jacqueroud L, Wierinckx A, et al. Snail family members unequally trigger EMT and thereby differ in their ability to promote the neoplastic transformation of mammary epithelial cells. PLoS One. 2014;9(3). doi:10.1371/journal.pone.0092254

53. Kim NH, Kim HS, Li XY, et al. A p53/miRNA-34 axis regulates Snail1-dependent cancer cell epithelial-mesenchymal transition. Journal of Cell Biology. 2011;195(3). doi:10.1083/jcb.201103097

